# Every individual makes a difference: A trinity derived from linking individual brain morphometry, connectivity and mentalising ability

**DOI:** 10.1101/2022.04.11.487870

**Authors:** Zhaoning Li, Qunxi Dong, Bin Hu, Haiyan Wu

**Affiliations:** Centre for Cognitive and Brain Sciences and Department of Psychology, University of Macau, Taipa, Macau SAR, China; School of Medical Technology, Beijing Institute of Technology, Beijing, China

**Keywords:** Mentalising, Surface-based multivariate morphometry statistics, Resting-state functional connectivity, Interactive mentalisation questionnaire, Inter-subject representational similarity analysis, Dyadic regression analysis

## Abstract

Mentalising ability, indexed as the ability to understand others’ beliefs, feelings, intentions, thoughts and traits, is a pivotal and fundamental component of human social cognition. However, considering the multifaceted nature of mentalising ability, little research has focused on characterising individual differences in different mentalising components. And even less research has been devoted to investigating how the variance in the structural and functional patterns of the amygdala and hippocampus, two vital subcortical regions of the ‘social brain’, are related to inter-individual variability in mentalising ability. Here, as a first step toward filling these gaps, we exploited inter-subject representational similarity analysis (IS-RSA) to assess relationships between amygdala and hippocampal morphometry (surface-based multivariate morphometry statistics, MMS), connectivity (resting-state functional connectivity, rs-FC) and mentalising ability (interactive mentalisation questionnaire (IMQ) scores) across the participants (*N* = 24). In IS-RSA, we proposed a novel pipeline, i.e., computing patching and pooling operations-based surface distance (CPP-SD), to obtain a decent representation for high-dimensional MMS data. On this basis, we found significant correlations (i.e., secondorder isomorphisms) between these three distinct modalities, indicating that a trinity existed in idiosyncratic patterns of brain morphometry, connectivity and mentalising ability. Notably, a region-related mentalising specificity emerged from these associations: self-self and self-other mentalisation are more related to the hippocampus, while other-self mentalisation shows a closer link with the amygdala. Furthermore, by utilising the dyadic regression analysis, we observed significant interactions such that subject pairs with similar morphometry had even greater mentalising similarity if they were also similar in rs-FC. Altogether, we demonstrated the feasibility and illustrated the promise of using IS-RSA to study individual differences, deepening our understanding of how individual brains give rise to their mentalising abilities.

---

> ‘But this general character is contained in every individual character; without individual character there can be no general character. If all individual character were removed, what general character would remain?’
>
> — (Mao Tse-tung — On Contradiction ^1^)

## 1. Introduction

Every individual makes a ‘difference’, and one’s mentalising ability or theory of mind (ToM) (Schaafsma et al., 2015), one of the core functions in human social life (Frith & Frith, 2006), underlies this significance and uniqueness. Such ability is fundamental to social understanding and social learning, and its dysfunction is associated with various social disorders, such as autism spectrum disorder (ASD) (Schuwerk et al., 2019; Hyatt et al., 2020) and social anxiety disorder (Washburn et al., 2016). Though most previous work has described mentalising ability as a whole (i.e., the ability to reason about and infer others’ mental states) (Frith & Frith, 2005; Adolphs, 2009), mentalising is a multifaceted concept that refers to multiple constructs involved in treating others and ourselves as social agents (Allen & Fonagy, 2008). Accordingly, the work with fine-grained dissection of the mentalising components will advance our understanding of mentalising during live social interaction (Wu et al., 2020b). Moreover, previous research on the inter-individual variability in mentalising ability has typically focused on neurological and psychiatric disorders (Stuss et al., 2001; Kerr et al., 2003; Richell et al., 2003; Snowden et al., 2003). While confirming the importance of mentalising in social life, substantial mentalising differences also exist in healthy adults. Yet, it is quite difficult to capture these differences behaviourally because of the ceiling effects observed on standard laboratory tasks (Koster-Hale & Saxe, 2013). Additionally, rare studies have been devoted to investigating sub-components of mentalising ability. In light of the newly proposed interactive mentalising theory (IMT) (Wu et al., 2020b) and its related interactive mentalising questionnaire (IMQ) (Wu et al., 2022), our study aimed to investigate the inter-individual variability in three different but interactive mentalising components: self-self mentalisation (SS), self-other mentalisation (SO) and other-self mentalisation (OS) (See more details about IMQ in Section 2.5).

More specifically, we sought to map the individual difference nature of different mentalising components to the brain. In the past decades, researchers have found numerous brain regions involved in mentalising, including the medial prefrontal cortex (mPFC), posterior superior temporal sulcus (pSTS) and temporoparietal junction (TPJ) (Frith & Frith, 2001; Siegal & Varley, 2002; Saxe & Kanwisher, 2003; Amodio & Frith, 2006; Schurz et al., 2014; Biervoye et al., 2016; Wu et al., 2020a). Recent work has extended the neural profile of mentalising from the single brain region to a widely distributed network of brain regions termed the ‘mentalising network’ (MTN) (Wang et al., 2021). However, two different but same vital subcortical regions of the ‘social brain’ (Bickart et al., 2014; Montagrin et al., 2018), i.e., the amygdala and hippocampus, which may also be the neural basis associated with individual differences in different mentalising components, have been less explored. In particular, the unasked question is whether inter-individual variability in the structural or functional patterns of the above two brain regions is associated with that in different mentalising components. Here, we used surface-based multivariate morphometry statistics (MMS) (Wang et al., 2010, 2011) and resting-state functional connectivity (rs-FC) to characterise amygdala and hippocampal structures and functions, respectively (Sections 2.3 and 2.4). Our goal was to take the first step toward interrogating whether individual differences in amygdala or hippocampal morphometry or connectivity reflect differences in mentalising ability of individuals.

We addressed this goal by using inter-subject representational similarity analysis (IS-RSA) (Nummenmaa et al., 2012; Feilong et al., 2018; Nguyen et al., 2019; van Baar et al., 2019; Chen et al., 2020; Finn et al., 2020) to link individual brain morphometry, connectivity and mentalising ability (Section 2.6). Specifically, based on the MRI and psychometric data collected from twenty-four participants (Sections 2.1 and 2.2), we computed each subject pair’s dissimilarity to construct inter-subject dissimilarity matrices (IDMs) for the above three modalities. In particular, we proposed a pipeline, i.e., computing patching and pooling operations-based surface distance (CPP-SD), for IDM construction of high-dimensional MMS data (Fig. 5). In the end, we compared these patterns (i.e., IDMs) to detect the shared structure of different modalities. We predicted that the levels of mentalising ability would correlate positively with the dissimilarity in amygdala and hippocampal morphometry and connectivity. Moreover, we also predicted that dissimilarity in functional and structural patterns would positively covary with each other. In general, the above predictions were rooted in our **Hypothesis 1**: Three distinct modalities will share one essence, i.e., there is a structure that exists in idiosyncratic patterns of brain morphometry, connectivity and mentalising ability, and we termed it as ‘trinity’.

Along with the trinity hypothesis, we further hypothesised that there will be a region-related specificity in associations among different mentalising components and amygdala or hippocampal MMS and rs-FC (**Hypothesis 2**). Specifically, mounting evidence indicates that both hippocampal function and structure are highly associated with meta-cognition (Chua et al., 2006; Moritz et al., 2006; Allen et al., 2017; Ren et al., 2018; Zou & Kwok, 2022). Meanwhile, two well-established theories: 1) relational integration theory (O’Keefe & Nadel, 1978) (born in Tolman’s seminal concept of the cognitive map (Tolman, 1948) and has been applied outside the original domain of physical space to social space (Rubin et al., 2014; Eichenbaum & Cohen, 2014; Tavares et al., 2015; Wang et al., 2015; Schafer & Schiller, 2018)) and 2) constructive memory theory (Schacter, 2012), all point to a crucial role of the hippocampus in how human construct, interact with and predict the intentions and actions of others (Laurita & Nathan Spreng, 2017). Ergo, we reasoned that hippocampal morphometry and connectivity would be more correlated with meta-cognition (self-self mentalisation) and perspective-taking (self-other mentalisation). While for other-self mentalisation (the ability to see ‘ourselves from the outside’ (Asen & Fonagy, 2012)), a higher-order mentalising component for complex social interactions, especially for trust or deception (Wu et al., 2022), we speculated that the amygdala would show a closer link to this component based on the indispensable role of the amygdala in developing and expressing interpersonal trust (Koscik & Tranel, 2011; Haas et al., 2015; Santos et al., 2016; Eskander et al., 2020).

Given that brain structure and function are highly related (Batista-García-Ramó & Fernández-Verdecia, 2018), the following interrelated hypothesis (**Hypothesis 3**) was that subject pairs with similar hippocampal MMS will have even greater self-self and self-other similarity if they are also similar in hippocampal rs-FC. In a similar vein, subject pairs with similar amygdala MMS will have even greater other-self similarity if they are also similar in amygdala rs-FC. To test this hypothesis and explore potential interaction effects, we analysed mentalising similarity across subject dyads using the mixed-effects dyadic regression model (Chen et al., 2017; van Baar et al., 2021) (Section 2.7).

## 2. Methods

### 2.1. Participants

We recruited thirty-one healthy right-handed participants (15 females, age: mean ± *SD* = 23.74 ± 4.02). They participated in this study via an online recruiting system. All participants filled out a screening form and were included in the study only if they confirmed they were not suffering from any significant medical or psychiatric illness, not using the medication, or not drinking or smoking daily. All the procedures involved followed the Declaration of Helsinki and were approved by the local ethics committee. To ensure the data quality of morphometry, we followed the literature (Worker et al., 2018; Dong et al., 2019) in visually examining T1-weighted images for motion artefact, wrap-around and grey/white contrast and thus excluded seven participants. Finally, we kept data from the other twenty-four participants (11 females, age: mean ± *SD* = 23.29 ± 2.95) in the following analysis.

### 2.2. Resting-state fMRI and structural MRI dataset

We utilised the resting-state fMRI (rsfMRI) and structural MRI data collected from the same participants. Specifically, MRI data was collected with a General Electric 3T scanner (GE Discovery MR750). The rsfMRI data was collected using gradient-echo by an echo-planar imaging (EPI) sequence. Slices were acquired in an interleaved order, and the data consisted of 200 whole-brain volumes (repetition time (TR) = 2,000 ms, echo time (TE) = 21 ms, flip angle = 90°, slice number = 42, slice thickness = 3.5 mm, matrix size = 64 × 64, field of view (FOV) = 200 mm and voxel size = 3.1 × 3.1 × 3.5 mm^3^). T1-weighted structural images were acquired using a 3D magnetisation-prepared rapid gradient-echo (MPRAGE) sequence (TR = 2,530 ms, TE = 2.34 ms, flip angle = 7°, FOV = 256 mm, slice number = 176, slice thickness = 1 mm, in-plane matrix resolution = 256 × 256, FOV = 256 mm and voxel size = 1 × 1 × 1 mm^3^). All participants underwent the T1 and resting-state fMRI scanning. This rsfMRI dataset was also used in our previous study (Pang et al., 2022).

### 2.3. MMS: Surface-based multivariate morphometry statistics

Surface-based multivariate morphometry statistics (MMS) was proposed for brain local structural analysis (Wang et al., 2010, 2011). Since then, studies have demonstrated that MMS has a larger effect size than volume, area and other similar morphometry measures (Wang et al., 2010; Dong et al., 2019, 2020; Wu et al., 2021). In this work, we calculated amygdala and hippocampal MMS from raw MR images by using the MRI processing pipeline used in the paper (Dong et al., 2019), as illustrated in Fig. 1. Specifically, each hippocampal or amygdala surface was parameterised into 15,000 vertices (Dong et al., 2019; Yao et al., 2020). And each vertex MMS has two kinds of morphometry features: radial distance (RD) (Pizer et al., 1999; Thompson et al., 2004) and multivariate tensor-based morphometry (mTBM) (Davatzikos, 1996; Thompson et al., 2000; Wang et al., 2010). The RD (a scalar) represents the thickness of the shape at each vertex to the medial axis, which reflects the surface differences along the surface normal directions (Wang et al., 2011). The medial axis is determined by the geometric centre of the isoparametric curve on the computed conformal grid (Wang et al., 2011). The axis is perpendicular to the isoparametric curve, so the thickness can be easily calculated as the Euclidean distance between the core and the vertex on the curve. The vertex mTBM (a 3 × 1 positive definite matrix) captures deformations within local surfaces, such as rotation, dilation and shears with surfaces perpendicular to RD (Shi et al., 2013; Dong et al., 2020). Since the surface of the hippocampus or amygdala in each brain hemisphere has 15,000 vertices, the feature dimensionality of both hippocampus or amygdala for each subject is 60,000 (15,000 × 4) eventually.

**Figure 1:**
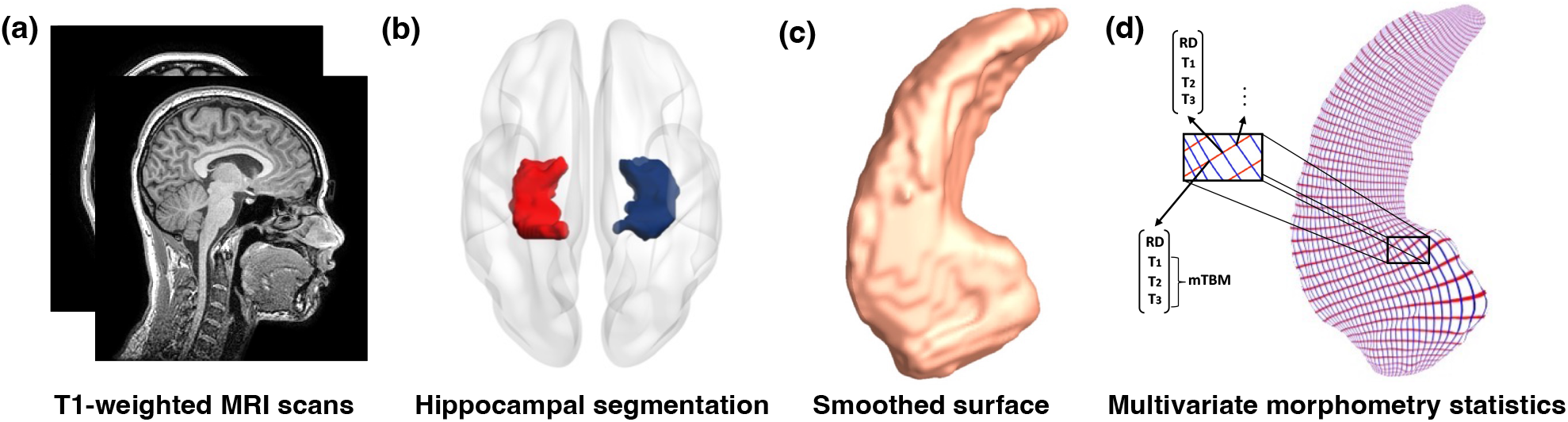
Processing pipeline of hippocampal morphometry data. (a-b) The hippocampal structures are segmented from registered T1-weighted MR images; (c) Smoothed hippocampal surfaces are then generated; (d) Surface-based multivariate morphometry statistics (MMS), including hippocampal radial distance (RD, a scalar) and the multivariate tensor-based morphometry (mTBM, a 3-dimensional vector) features at each vertex, are calculated after the surface parameterisation and fluid registration. Notice that we also applied the same pipeline to get amygdala morphometry data. Sub-figure (a) and (b) were visualised by using Nilearn (Abraham et al., 2014) and BrainNet Viewer (Xia et al., 2013), respectively.

### 2.4 Rs-FC: Resting-state functional connectivity

To profile amygdala and hippocampal functions, we utilised resting-state functional connectivity (rs-FC), which could predict social behaviours and thus has advanced our understanding of complex mental states (Eickhoff & Müller, 2015; Finn et al., 2015; Pezzulo et al., 2021; Pang et al., 2022). The rsfMRI preprocessing was adopted from our previous study (Pang et al., 2022). Specifically, the DPARSF V5.1 module (Yan et al., 2016) was used to preprocess rsfMRI data. The initial ten volumes were discarded because of instability in the MRI signal. Then, all slices were corrected for varying acquisition times based on each participant’s image series. T1-weighted MPRAGE structural images were aligned with the mean functional image after realignment. The structural images were segmented into grey matter, white matter and cerebrospinal fluid (Ashburner & Friston, 2005). To eliminate nuisance signals, the Friston 24-parameter model (Friston et al., 1996) was applied to remove head motion, the mean white matter and cerebrospinal fluid signals. The functional data from individual native space was transformed to the standard Montreal Neurological Institute space ^2^, and then spatial smoothing (FWMH kernel: 6 mm) was applied for connectivity analysis. Additionally, temporal filtering (0.01-0.1 Hz) was applied to the time series. Then, to conduct the rs-FC analysis, we extracted the BOLD time series of 116 regions in the whole brain according to the Automated Anatomical Labelling (AAL) 116 atlas (Tzourio-Mazoyer et al., 2002). Then, we obtained a 116 × 116 functional connectivity (FC) matrix by Pearson’s correlation analysis between the averaged BOLD time series of each pair of brain regions. We calculated the static FC matrix subject by subject, resulting in 24 FC matrices. Notably, with our specific research interest in the amygdala and hippocampus in this study, we only used FC between bilateral amygdala or hippocampus and other brain regions from the AAL 116 atlas for the following analysis (Fig. 2 shows their group-average profiles).

**Figure 2:**
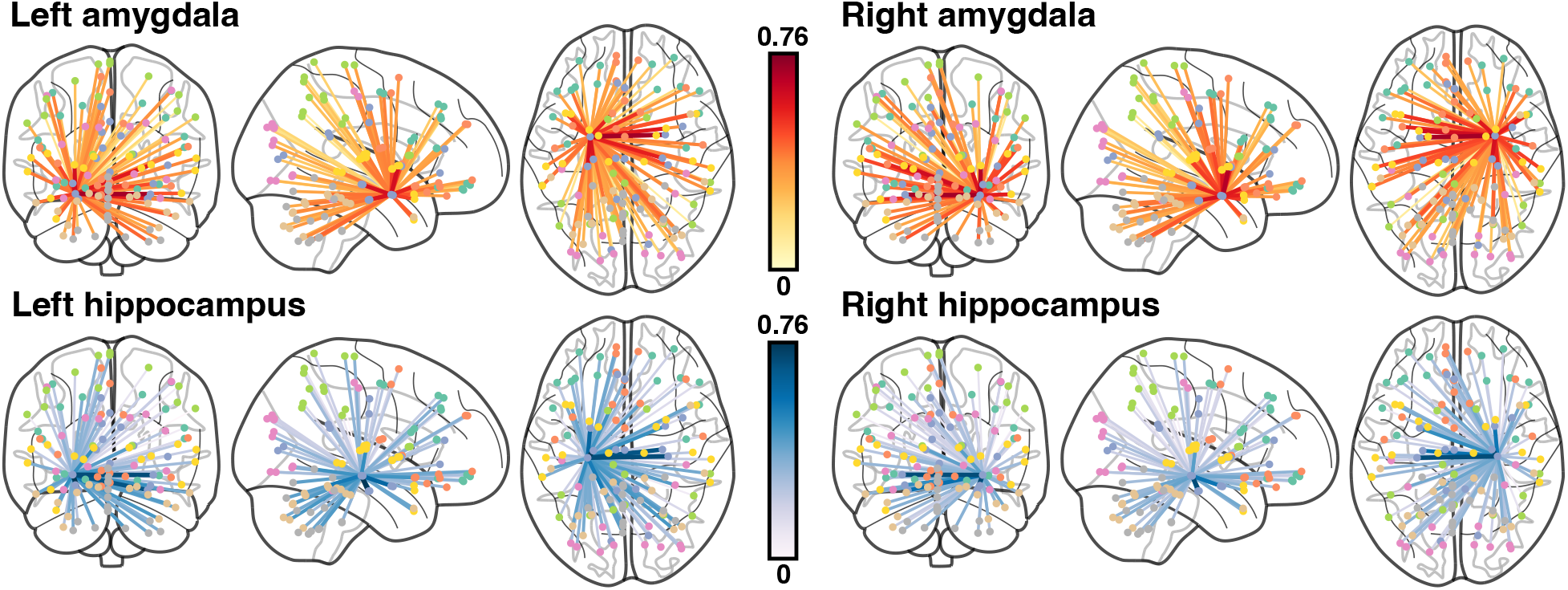
Group-average profiles of resting-state functional connectivity between seeds (top: amygdala, bottom: hippocampus) and other brain regions from the AAL 116 atlas, separated by hemisphere. The figure was visualised using Nilearn (Abraham et al., 2014).

### 2.5. IMQ: Interactive mentalisation questionnaire

While mentalising ability has classically been studied as a whole capacity (Frith & Frith, 2005; Adolphs, 2009), different types of mentalising may occur in social interaction, and it requires measures of the mentalising components, as people’s inner states are not directly observable (Wu et al., 2020b). Recently, the newly developed interactive mentalising questionnaire (IMQ) (Wu et al., 2022) offers us opportunities to capture different but interactive mentalising components, and thus we could further focus on their neural association related to key brain regions in social interaction. The IMQ contains 20 items and covers three mentalising components, i.e., self-self mentalisation (SS, the ability to look inward to self-monitor and assess thought processes), self-other mentalisation (SO, the ability to infer the mental states and thoughts of others) and other-self mentalisation (OS, the ability to make inferences about how much insight one think other agents have into one’s own thoughts and intentions). In this study, participants completed IMQ after the scanning session, and we visualised their position in a 3-dimensional space (composed of scalar summary scores of SS, SO or OS) based on their IMQ scores in Fig. 3.

**Figure 3:**
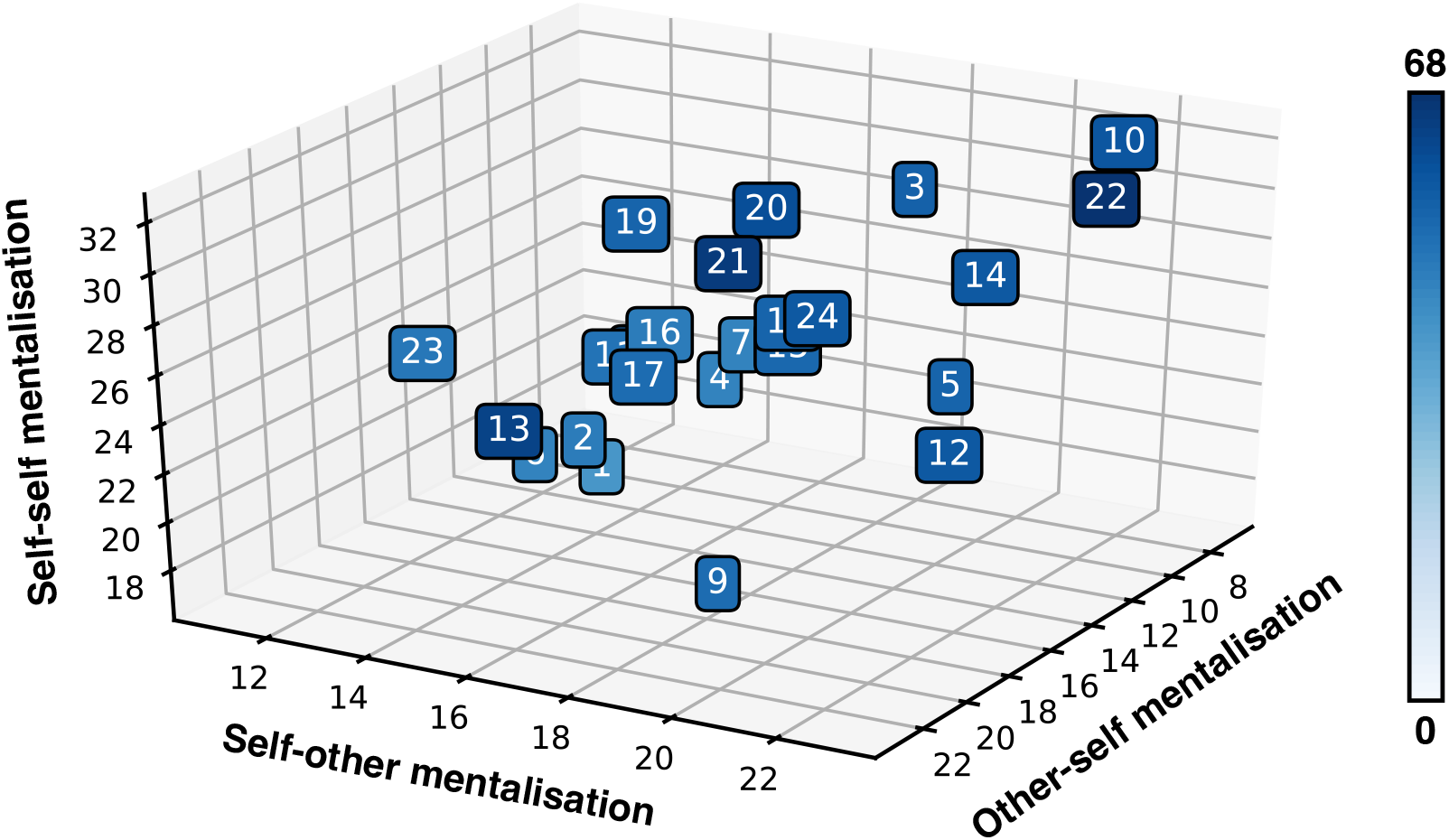
Visualisation of IMQ scores of different participants. We visualised the position of each participant based on their IMQ scores, which can be divided into three parts: self-self mentalisation (SS), self-other mentalisation (SO) and other-self mentalisation (OS). The colour intensity denotes the participants’ overall summary score for the IMQ. The figure was visualised by using Matplotlib (Hunter, 2007).

### 2.6. IS-RSA: Inter-subject representational similarity analysis

We exploited inter-subject representational similarity analysis (IS-RSA) (Nummenmaa et al., 2012; Feilong et al., 2018; Nguyen et al., 2019; van Baar et al., 2019; Chen et al., 2020; Finn et al., 2020) to link individual brain morphometry, connectivity and mentalising ability (i.e., test **Hypothesis 1**), as shown in Fig. 4. This analytic technique enables us to explore how individual differences in one modality (e.g., rsfMRI) is related to that in another modality (e.g., behavioural disposition) using second-order isomorphism (Shepard & Chipman, 1970) akin to representational similarity analysis (RSA) (Kriegeskorte et al., 2008; Kriegeskorte & Kievit, 2013; Popal et al., 2019). The intuition is that if individuals are more similar in one modality (e.g., MMS), they will also exhibit a higher similarity in another modality (e.g., IMQ scores). We operationalised this by computing three types of inter-subject dissimilarity matrices (IDMs), which characterise related modality’s representations in terms of subject-to-subject differences for all the MMS, rs-FC and IMQ. Then we can compute the similarity among these modalities by detecting the degree to which their IDMs agree. By directly comparing the representational geometry (Kriegeskorte et al., 2008; Kriegeskorte & Kievit, 2013) on the level of IDMs, we circumvent the problem of defining sophisticated mapping functions between different modalities.

**Figure 4:**
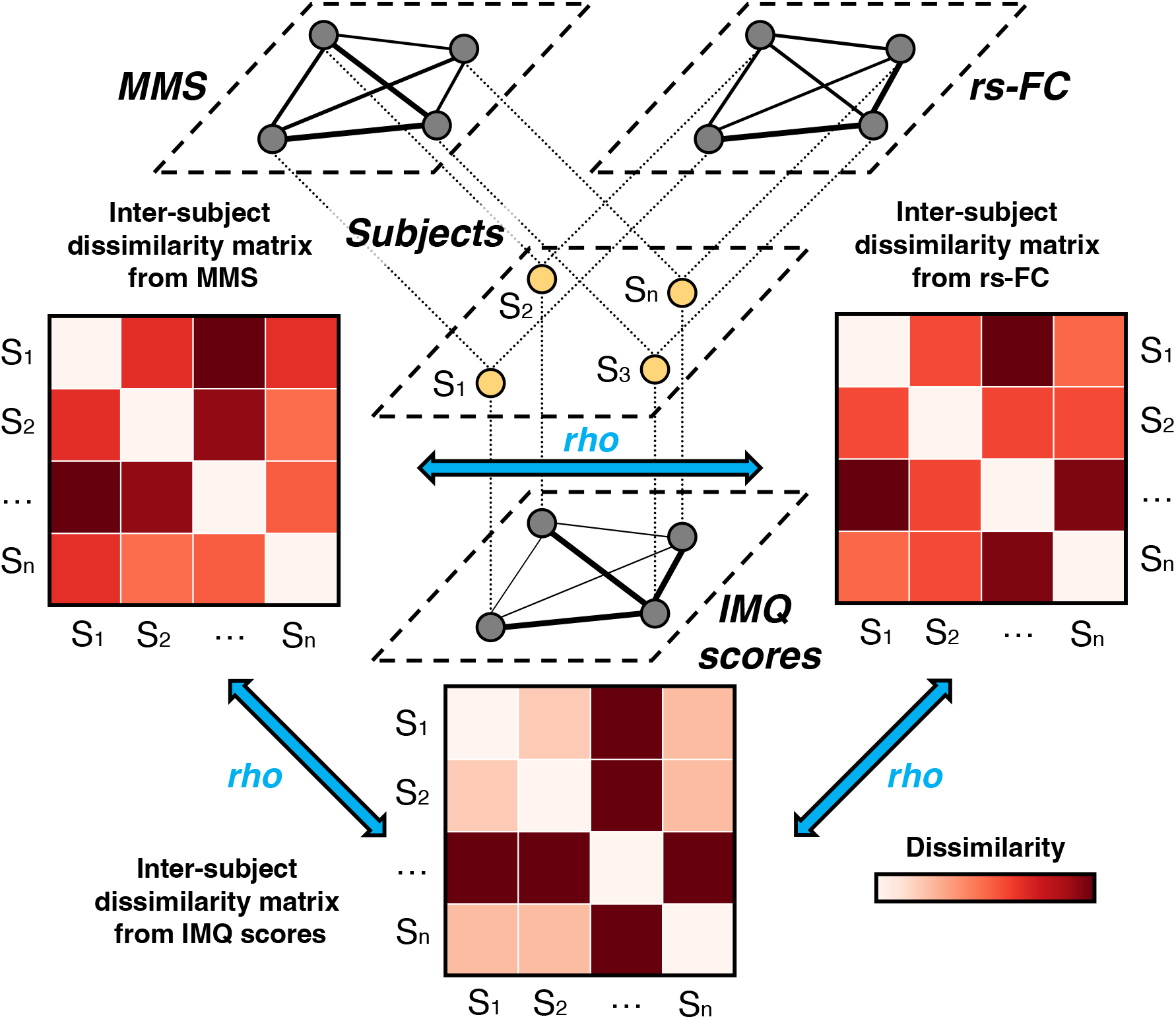
A schematic illustration of the inter-subject representational similarity analysis framework. For each subject (middle layer), we can compute the dissimilarity between each subject pair by using IMQ scores (bottom ‘IMQ scores’ layer), the pattern of MMS (top-left ‘MMS’ layer) and the pattern of rs-FC (top-right ‘rs-FC’ layer) for their amygdala and hippocampus. The top and bottom layers depict weighted graphs using adjacency matrices, i.e., inter-subject dissimilarity matrices, in which thicker lines denote increased dissimilarity between subjects. In IS-RSA, we constructed inter-subject dissimilarity matrices for all the rs-FC, MMS and IMQ scores and then compared them using Spearman rho rank-order correlation. In this way, we can detect shared structure between amygdala and hippocampal morphometry, connectivity and mentalising components.

**Figure 5:**
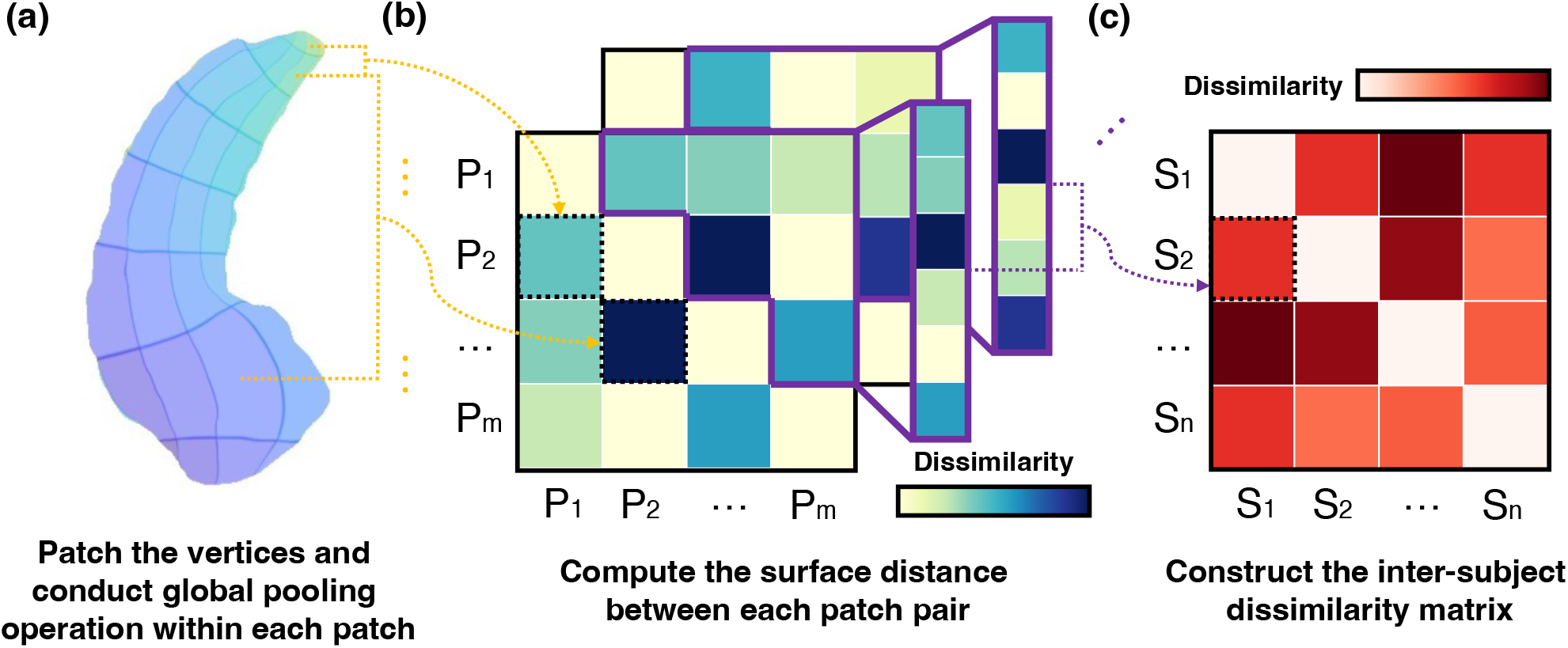
The pipeline of constructing inter-subject dissimilarity matrix for hippocampal MMS data. (a) We first patch the vertices on each hippocampal surface and conduct global pooling operation (Collobert et al., 2011) within each patch to obtain the vector representation for each path; (b) Then we compute the surface distance between each patch pair for the whole surface, resulting in an m-dimensional symmetric matrix (m denotes the patch number); (c) We extract the upper (or lower) off-diagonal triangle of this matrix and reshape it to a vector as the representation of one’s hippocampal morphometry, and thus we could compute the dissimilarity between each subject pair via different distance metrics. Notice that we also applied the same construction pipeline to amygdala MMS data.

Moving closer to individual differences in brain morphometry, we computed MMS IDMs separately from amygdala or hippocampal surface (one 15,000 × 4 surface for each brain region). Specifically, we proposed a pipeline, i.e., computing patching and pooling operations-based surface distance (CPP-SD), to obtain a decent surface representation for high-dimensional MMS data. Then, we computed pairwise distances again, not based on original high-dimensional MMS data but on the extracted surface distances matrices, as depicted in Fig. 5. We speculated that constructing MMS IDM using CPP-SD would improve the IDM correspondences compared to the conventional way, i.e., basing IDM directly on reshaped surfaces (a single vector of 60,000 values for each surface), which may treat noisy values as equally important to values that carry the signal (Kaniuth & Hebart, 2022). To construct rs-FC IDM separately for the amygdala and hippocampus, we computed the dissimilarity between pairs of subjects’ amygdala or hippocampal rs-FC (one vector of 116 values for each brain region). In the same way, we constructed IMQ IDM separately for each mentalising component (i.e., SS, SO and OS).

Finally, we quantified representational similarity as the Spearman rho rank-order correlation between the upper (or lower) off-diagonal triangles of IDMs. We had four MMS IDMs, four rs-FC IDMs (for bilateral amygdala and hippocampus, separately) and three IMQ IDMs (for three mentalising components), so this procedure yielded 40 representational similarity values in our analysis. To assess statistical significance, we followed the literature (Bonner & Epstein, 2018; Lu & Ku, 2020) in using a permutation test, in which the rows and columns of one of the IDMs were randomly shuffled, and a Spearman correlation between IDMs was calculated over 10,000 iterations. Then we calculated *p*-values from this permutation distribution for a one-tailed test as follows:

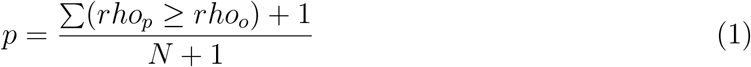

where *rho_p_* and *rho_o_* denote the Spearman correlations from the permutation distribution and the original data, respectively. We thresholded all permuted *p*-values at a false-discovery rate (FDR) of 0.05 (Benjamini–Hochberg method (Benjamini & Hochberg, 1995)).

During the construction of IDMs, different types of parameters might affect the final correspondence. Empirically, we tried different patch sizes, including 5 × 5, 10 × 10, 25 × 25 and 50 × 50 vertices, and different pooling operations, including max-, mean-, min-over-vertex operation or a combination of these three kinds of operations for constructing MMS IDM. We also tried different distance metrics, including Pearson distance, Euclidean distance, Mahalanobis distance, cosine distance and Manhattan distance, for constructing all three types of IDMs, plus word mover’s distance (Kusner et al., 2015) and word rotator’s distance (Yokoi et al., 2020), for constructing MMS IDM (with patching and pooling operations but without computing surface distance in this case). To ensure no single IMQ item could drive the dissimilarity between subjects when applying the above distance metrics, we followed the literature (Chen et al., 2020) in normalising IMQ items to the range [0, 1] prior to computing pairwise multivariate distances. Besides these distance metrics, we also tried absolute distance, Anna Karenina distances (Finn et al., 2020) (all high scorers are alike, and all low scorers are low-scoring in their own way), including mean distance, minimum distance and the product of the absolute and minimum distance, and reversed Anna Karenina distance (Finn et al., 2020) (all low scorers are alike, all high scorers are high-scoring in their own way), i.e., maximum distance, for scalar summary scores of mentalising components.

We performed a grid search (Feurer & Hutter, 2019) over the above parameter space and selected the optimal parameter configuration with the highest average IDM correspondence from subject-wise bootstrapping (Chen et al., 2016). Specifically, in each of the 1,000 repetitions, we performed resampling with replacement from the 24 original individuals. This operation would involve non-informative dissimilarity values of the IDM diagonals (comparisons of subjects to themselves), so we excluded these values (less than 4.2%) to avoid overestimating representational similarity values (Kriegeskorte et al., 2008). Then we computed IDMs based on the bootstrapped samples and obtained average IDM correspondence and 95% confidence intervals accordingly.

### 2.7. Dyadic regression analysis

To evaluate **Hypothesis 3** about the interaction between MMS and rs-FC similarity in predicting mentalising similarity ^3^, we further conducted the dyadic regression analysis (Chen et al., 2017; van Baar et al., 2021), i.e., a regression analysis in which each observation portrays the similarity between a subject pair. By using linear mixed-effects modelling, this analysis could account for inherent statistical dependencies between all possible subject pairs (dyads) (Chen et al., 2017; Parkinson et al., 2018) and evaluate the interactions between MMS and rs-FC simultaneously, which is impossible in standard RSA. Here, we used Pymer4 (Jolly, 2018) to implement the mixed-effects regression model, which included a random subject intercept for both subjects involved in that subject pair. The model thus followed the formulation as shown below:

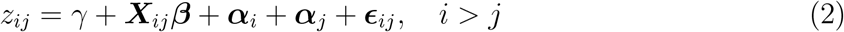

where *i* and *j* denote two subjects in a dyad, and *z_ij_* represents pairwise observations (i.e., the similarity between two subjects in one mentalising component). We regressed *z_ij_* onto fixed effects ***X**_ij_* (i.e., MMS similarity, rs-FC similarity and their interaction), random effects ***α**_i_* and ***α**_j_* (***α*** ~ ***N*** (**0**, ***σ_α_***)) attributable to subjects *i* and *j*, and the residual term ***ϵ**_ij_* (***ϵ*** ~ ***N*** (**0**, ***σ_ϵ_***)) for the dyad (*i, j*).

## 3. Results

### 3.1. Results of IS-RSA

We performed inter-subject representational similarity analysis (IS-RSA) to test **Hypothesis 1**, i.e., whether a trinity could derive from linking individual brain morphometry, connectivity and mentalising ability. In other words, we expected to observe positive correlations among these three seemingly distinct modalities. Consistent with our **Hypothesis 1**, the results (see Fig. 6 and Table S1) revealed that a trinity indeed existed in idiosyncratic patterns with all correlations significantly greater than zero (all the *p*-values < .05 after applying the FDR correction for multiple comparisons), indicating brain morphometry, connectivity and mentalising ability were fundamentally related.

**Figure 6:**
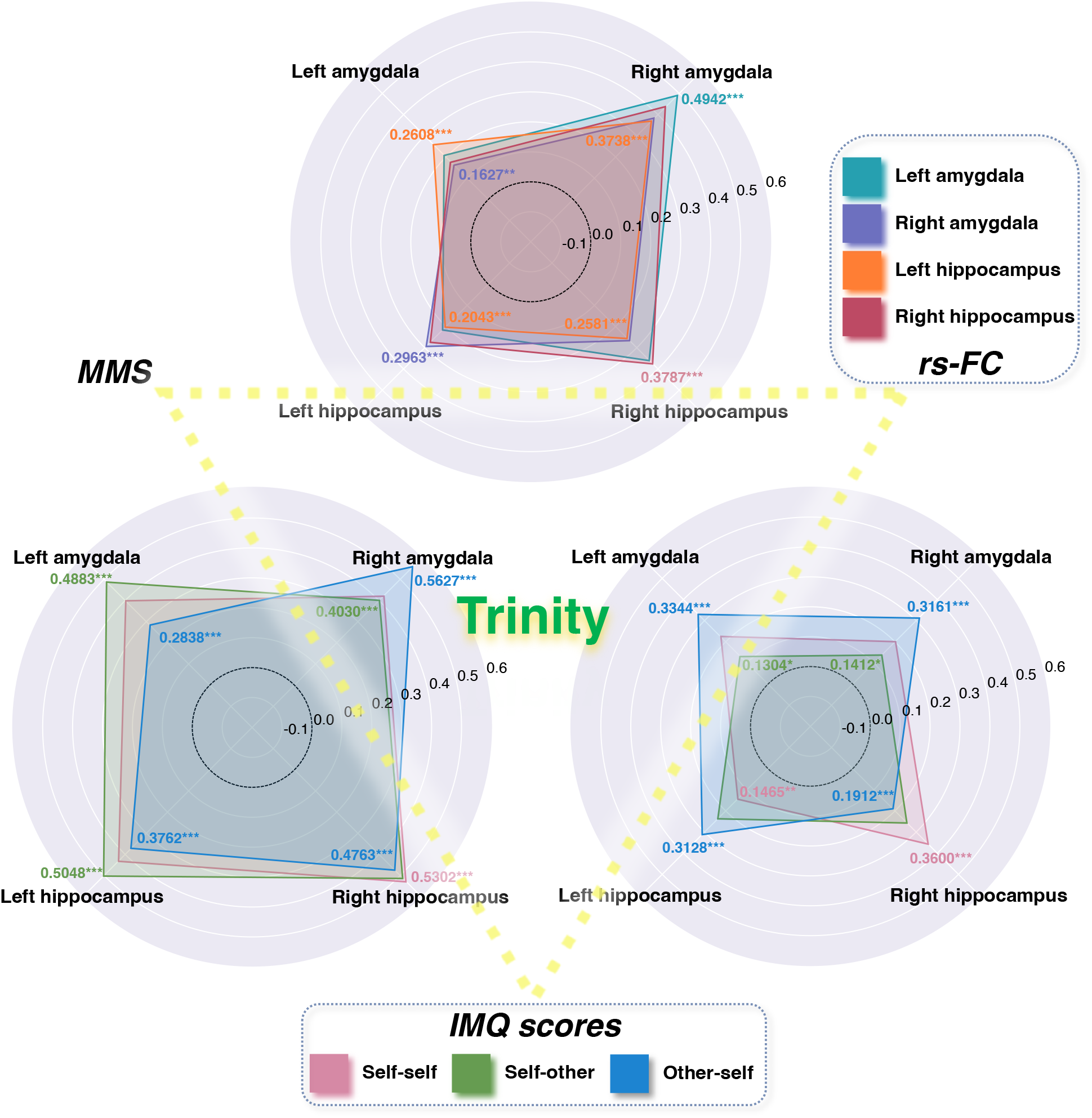
A trinity existed in idiosyncratic patterns of brain morphometry, connectivity and mentalising ability. The bottom left, bottom right and top radar charts depict representational similarity (in Spearman rho rank-order correlation units) between IDMs of IMQ scores (including three different components) and MMS (including bilateral amygdala and hippocampus), IMQ scores and rs-FC (including bilateral amygdala and hippocampus), and rs-FC and MMS. We used the FDR correction for multiple comparisons across all 40 statistical tests (12 for IMQ-MMS, 12 for IMQ-rs-FC and 16 for MMS-rs-FC). All correlations were significantly greater than zero (all corrected *p*-values < .05). Detailed results were reported in Table S1. * *p* < .05, ** *p* < .01, *** *p* < .001.

In particular, when comparing similarities between IMQ scores and MMS, the highest similarities were found in the right hippocampus (SS: *rho* = 0.5302, *p* < .001; SO: *rho* = 0.5156, *p* < .001) and the right amygdala (OS: *rho* = 0.5627, *p* < .001) (Table S1a). When comparing similarities between IMQ scores and rs-FC, the highest similarities were found in the right hippocampus (SS: *rho* = 0.3600, *p* < .001; SO: *rho* = 0.2580, *p* < .001) and the left amygdala (OS: *rho* = 0.3344, *p* < .001) (Table S1b). When comparing similarities between rs-FC and MMS, the highest similarities were found in the right amygdala (rs-FC_*LA*_: *rho* = 0.4942, *p* < .001; rs-FC_*RA*_: *rho* = 0.3848, *p* < .001; rs-FC_*RA*_: *rho* = 0.3738, *p* < .001; rs-FC_*RA*_: *rho* = 0.4421, *p* < .001) (Table S1c).

After finding support for **Hypothesis 1**, we performed two-sided Wilcoxon signed-rank tests based on the bootstrapped samples to further test **Hypothesis 2**, i.e., (a) whether an IDM derived from either SS or SO is more related to an IDM derived from hippocampal MMS or rs-FC compared to that derived from amygdala MMS or rs-FC; (b) whether an IDM derived from OS is more related to an IDM derived from amygdala MMS or rs-FC compared to that derived from hippocampal MMS or rs-FC. We corrected multiple comparisons by controlling the expected FDR at 0.05. As shown in Table 1, we found that the IDM derived from the right hippocampal MMS or rs-FC was, on average, significantly more correlated with that from SS (MMS_*RH*_ : *rho_mean_* = 0.5168, rs-FC_*RH*_ : *rho_mean_* = 0.3434) or SO (rs-FC_*RH*_ : *rho_mean_* = 0.2427). Although the average IDM correspondence between the right hippocampal MMS and SO was not significantly stronger than that between the left hippocampal or left amygdala MMS and SO, it was the highest correlation score derived from quantifying representational similarity between MMS and SO (MMS_*RH*_ : *rho_mean_* = 0.4766). We also found that the average IDM correspondence between the right amygdala MMS or the left amygdala rs-FC and OS was significantly stronger than that between the other brain regions of interest and OS (MMS_*RA*_: *rho_mean_* = 0.5153, rs-FC_*LA*_: *rho_mean_* = 0.3164). In summary, the results corroborated our proposed **Hypothesis 2**.

**Table 1:**
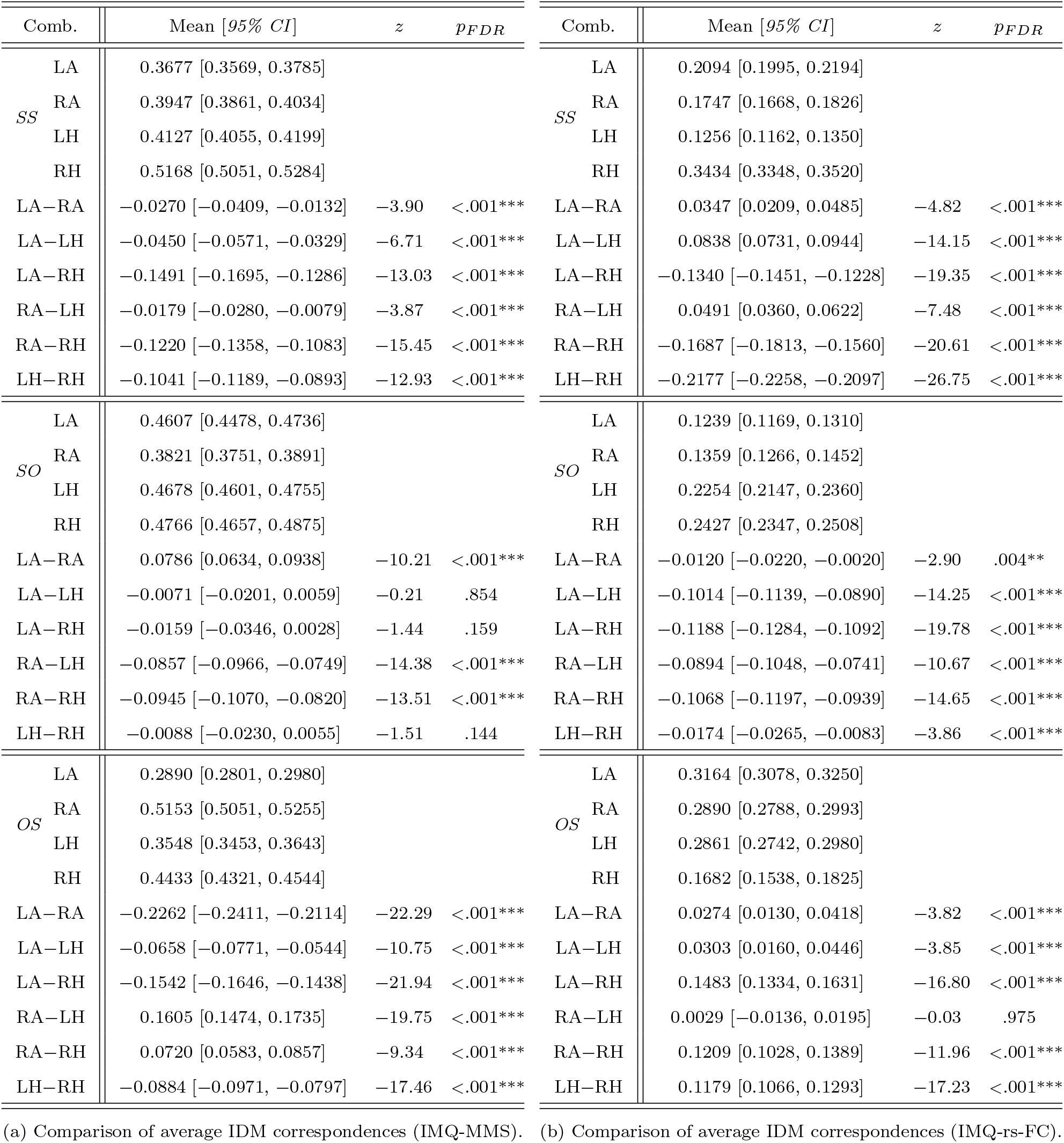
Comparison of average IDM correspondences from bootstrapping among bilateral amygdala and hippocampus. ‘Comb.’ for combinations; ‘LA’ for the left amygdala; ‘RA’ for the right amygdala; ‘LH’ for the left hippocampus; ‘RH’ for the right hippocampus. Mean values and 95% confidence intervals (numbers in parentheses) were obtained by bootstrapping subjects. All the *z*-values and *p*-values were derived from two-sided Wilcoxon signed-rank tests. We used FDR correction for multiple comparisons across all 36 statistical tests (18 for IMQ-MMS and 18 for IMQ-rs-FC). * *p* < .05, ** *p* < .01, *** *p* < .001.

### 3.2. Results of dyadic regression analysis

After finding support for our **Hypotheses 1** and **2**, we then turned to **Hypothesis 3** that subject pairs with similar hippocampal (or amygdala) morphometry will have even greater SS and SO similarity (or OS similarity) if they are also similar in hippocampal (or amygdala) rs-FC. For the dyadic regression analysis, we chose the right hippocampus and bilateral amygdala from **Hypothesis 2** as brain regions of interest. We noted that all regressors were only weakly correlated (*rho* < |0.28|), ensuring estimable linear mixed-effects models. The results first revealed that the interaction between MMS and rs-FC similarity drove the similarities of all three mentalising components (see Fig. 7a). This effect was significant over the right hippocampus in predicting SS similarity (*β* = 0.159, *SE* = 0.019, *p* < .001) and SO similarity (*β* = 0.050, *SE* = 0.021, *p* = .020) and left amygdala in predicting OS similarity (*β* = 0.046, *SE* = 0.023, *p* = .046) (Table 2).

**Table 2:** MMS and rs-FC similarity of the hippocampus and amygdala (right hippocampus, RH, left amygdala, LA, and right amygdala, RA) interacted to drive the similarities of all three mentalising components, as shown in Fig. 7. Each model included crossed random effects for subjects and fixed effects for MMS similarity (z-scored), rs-FC similarity (z-scored) and their interaction, with mentalising similarity (z-scored) as the outcome variable. ‘Comb.’ for combinations. * *p* < .05, ** *p* < .01, *** *p* < .001.

| Comb. | Term | $\beta$ | $SE$ | $t$ | $df$ | $p$ |
| --- | --- | --- | --- | --- | --- | --- |
| SS-RH | MMS similarity | 0.272 | 0.048 | 5.62 | 494.2 | <.001*** |
|  | rs-FC similarity | 0.057 | 0.026 | 2.22 | 525.8 | .027* |
| | MMS similarity ×<br>rs-FC similarity | 0.159 | 0.019 | 8.23 | 502.4 | <.001*** |
| SO-RH | MMS similarity | 0.084 | 0.031 | 2.70 | 532.3 | <.007** |
|  | rs-FC similarity | 0.002 | 0.028 | 0.07 | 524.3 | .944 |
| | MMS similarity ×<br>rs-FC similarity | 0.050 | 0.021 | 2.33 | 505.7 | .020* |
| OS-LA | MMS similarity | 0.181 | 0.040 | 4.52 | 547.6 | <.001*** |
|  | rs-FC similarity | 0.010 | 0.031 | 0.32 | 529.1 | .746 |
| | MMS similarity ×<br>rs-FC similarity | 0.046 | 0.023 | 2.00 | 508.2 | .046* |
| OS-RA | MMS similarity | -0.009 | 0.034 | -0.27 | 538.0 | .789 |
|  | rs-FC similarity | 0.055 | 0.025 | 2.18 | 515.8 | .030* |
| | MMS similarity ×<br>rs-FC similarity | -0.033 | 0.020 | -1.68 | 506.0 | .094 |

**Figure 7:**
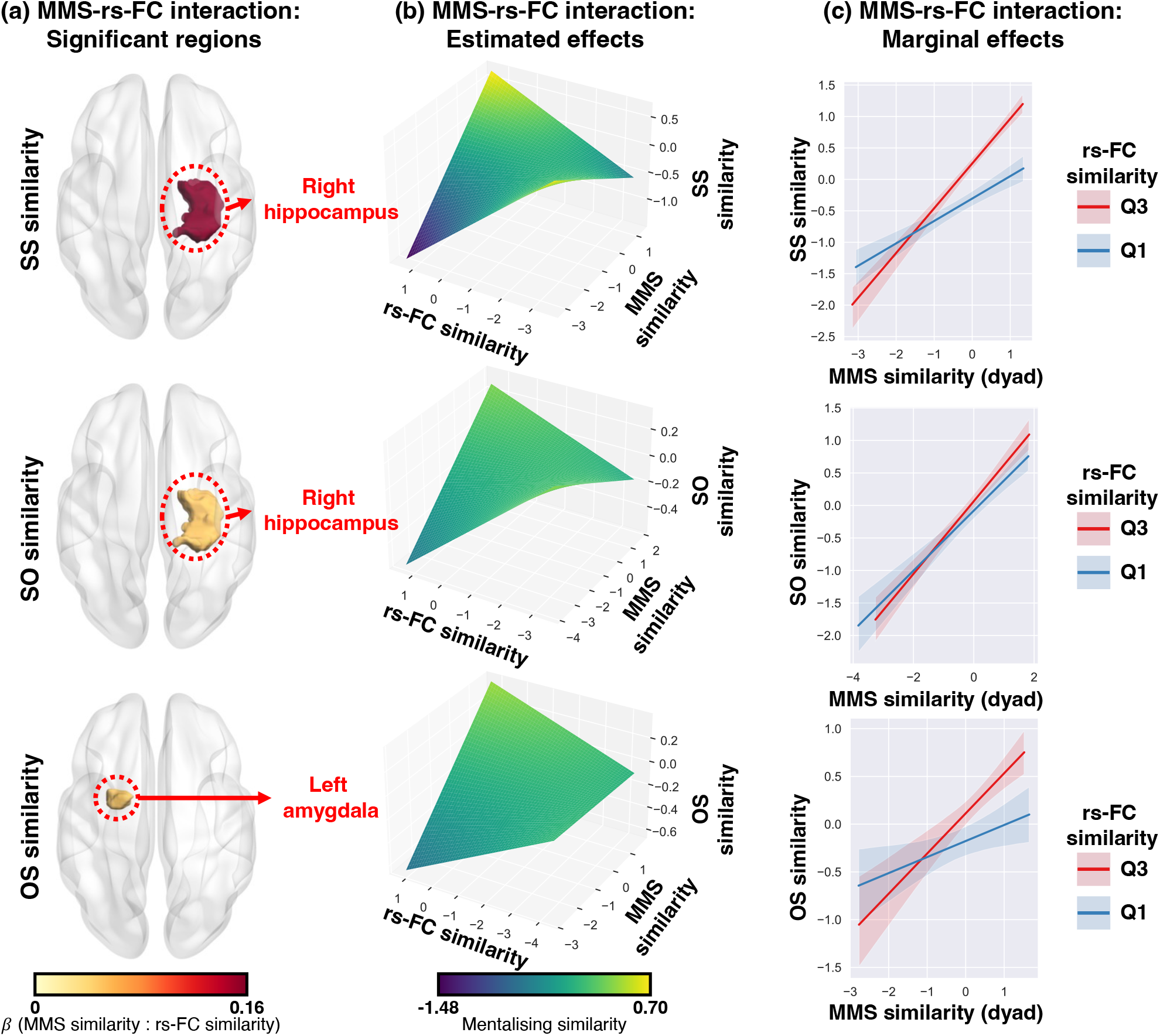
(a) The interaction between MMS and rs-FC similarity drove the similarities of all three mentalising components; (b) Simulating mentalising similarity from the regression model revealed that rs-FC similarity amplified mentalising similarity: MMS similarity more strongly predicted mentalising similarity in subject pairs with similar rs-FC (high-rs-FC similarity); (c) Estimated marginal effects for mentalising similarity by rs-FC similarity (lower quartile, Q1, and upper quartile, Q3) confirmed this interpretation: High-rs-FC similarity boosted mentalising similarity in subject pairs with high-MMS similarity and lowered mentalising similarity for dyads with low-MMS similarity. Shaded areas denote 95% confidence intervals; Sub-figure (a), (b) and (c) were visualised by using BrainNet Viewer (Xia et al., 2013), Matplotlib (Hunter, 2007) and seaborn (Waskom, 2021) respectively. Regression coefficients were reported in Table 2.

More importantly, to understand the directionality of the observed MMS-rs-FC interaction effects, we plotted the mentalising similarity predicted by the fitted regression models in the significant regions, including the right hippocampus and left amygdala (see Fig. 7b, Fig. 7c and Table 2). Consistent with our hypothesis, high-rs-FC similarity amplified mentalising similarity, such that two individuals who were both similar in right hippocampal (or left amygdala) rs-FC and MMS would have significantly higher similarity in self-self and self-other mentalisation (or other-self mentalisation).

### 3.3. Results of pipelines for constructing inter-subject MMS dissimilarity matrix

To evaluate whether our proposed pipeline CPP-SD, i.e., computing patching and pooling operations-based surface distance, could increase the correspondence between MMS IDM and the other IDM, we compared CPP-SD with pipeline 2 (with patching and pooling operations but without computing surface distance) and pipeline 1 (without patching and pooling operations and computing surface distance). Fig. 8 shows the comparison of average IDM correspondences from bootstrapping among these three pipelines. Specifically, we found that pipeline 2-based IS-RSA reveals a significantly stronger IDM correspondence than pipeline 1-based IS-RSA for 27 cases (96.43%) (two-sided FDR-corrected Wilcoxon signed-rank tests, similarly hereinafter). Only in 1 case (3.57%, target IDM: rs-FC) the difference between pipeline 2-based IS-RSA and pipeline 1-based IS-RSA was not significant, indicating the effectiveness of patching and pooling operations. Compared with pipeline 2, we also found that basing IS-RSA on CPP-SD further significantly improved the IDM correspondence for 22 cases (78.57%), with 3 cases (10.71%, target IDM: IMQ scores) that showed significantly worse performance of IS-RSA and 3 cases (10.71%, target IDM: rs-FC) with a non-significant difference. Together, these results demonstrated that our proposed CPP-SD robustly improved the correspondence between MMS and the other two modalities.

**Figure 8:**
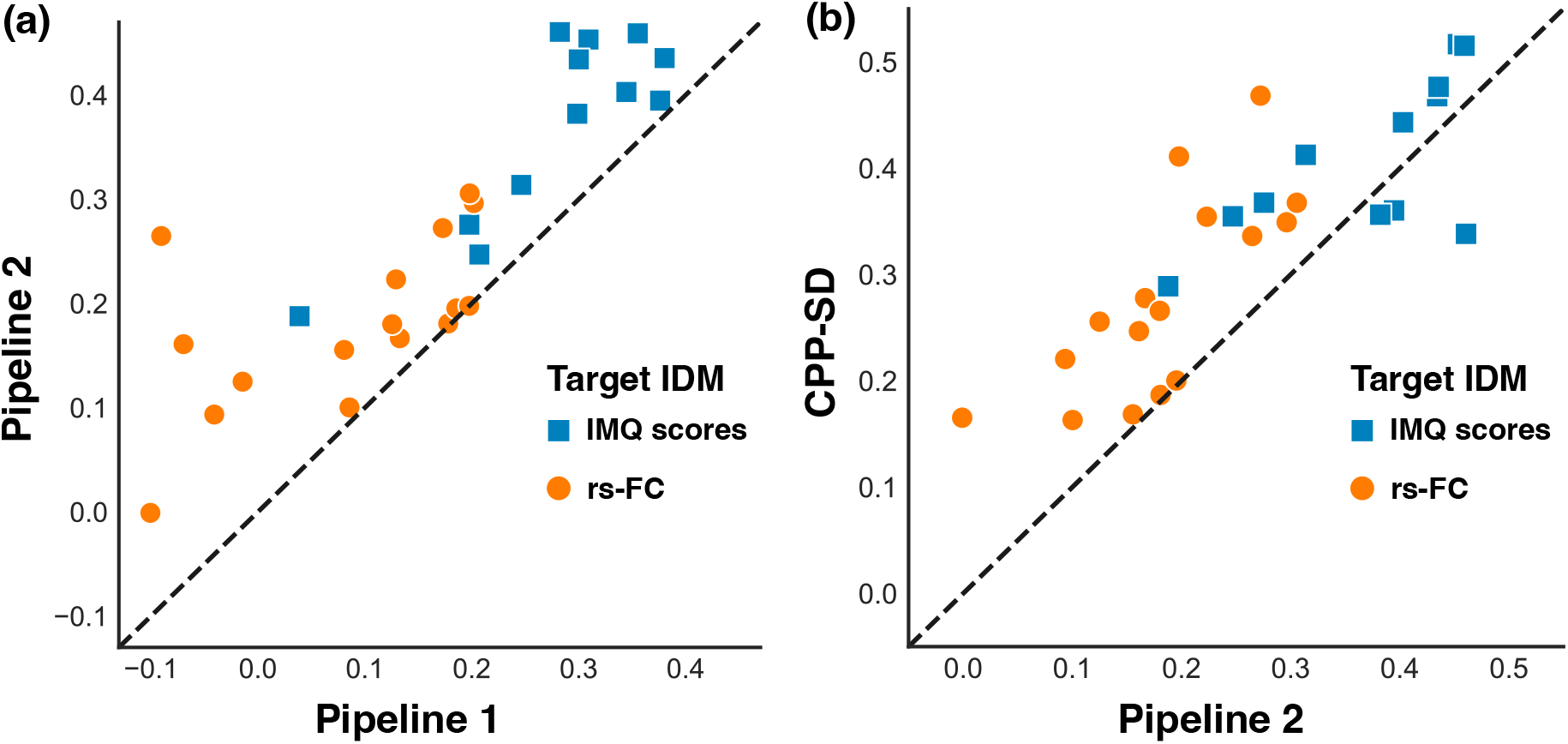
Comparison of average IDM correspondences from bootstrapping among three pipelines. (a) Pipeline 2-based (with patching and pooling operations but without computing surface distance) IS-RSA reveals a stronger IDM correspondence than pipeline 1-based (without patching and pooling operations and computing surface distance) IS-RSA for all 28 comparisons. Among them, pipeline 2-based IS-RSA significantly out-performed pipeline 1-based IS-RSA in 27 cases, usually leading to increases in the representational similarity between two IDMs; (b) For most comparisons, basing IS-RSA on CPP-SD (computing patching and pooling operations-based surface distance) further improved the IDM correspondence. Of all 28 comparisons, CPP-SD-based IS-RSA significantly outperformed pipeline 2-based IS-RSA in 22 cases, often leading to increases in the representational similarity between two IDMs. The figure was visualised using seaborn (Waskom, 2021).

## 4. Discussion

Social behaviours, to some degree, largely rely on various mentalising components (Wu et al., 2020b). However, very little is known about how variation in these components (i.e., self-self mentalisation, self-other mentalisation and other-self mentalisation) mapped onto individual variation in the amygdala and hippocampus, two vital subcortical regions of both the traditionally ‘mammalian brain’ (MacLean, 1990) and newly developed ‘social brain’ (Bickart et al., 2014; Montagrin et al., 2018). As a first step toward filling this gap, the goal of our study was to provide a thorough investigation of individual differences in mentalising components, amygdala and hippocampal morphometry and connectivity by detecting shared structure among all three using inter-subject representational analysis (IS-RSA). Consistent with our **Hypothesis 1**, we found that a trinity existed in idiosyncratic patterns of amygdala and hippocampal morphometry, connectivity and mentalising ability (Fig. 6), advancing our understanding of the neural basis of mentalising. In this regard, our study also served as a proof of principle for using IS-RSA to directly link (i.e., bypassing the problem of defining sophisticated mapping functions between different modalities) idiosyncratic patterns of the above three modalities. Our IS-RSA-based trinity framework could easily transfer to other brain regions and is straightforward to implement. In this respect, we believe that it is a powerful starship by which we could go where no one has gone before and specifically move from asking simple questions about mentalising ability to asking how individual variations emerge from complex social interactions and what neurocognitive mechanisms underlie these variations.

In line with our **Hypothesis 2**, we also obtained a region-related specificity in associations among mentalising components and amygdala or hippocampal MMS and rs-FC (Table 1a and 1b). The specificity is in three aspects. First, the variation of self-self mentalisation, i.e., metacognition, exhibited greater representational similarity to individual variation in hippocampal MMS or rs-FC relative to amygdala MMS or rs-FC, indicating a close association between the meta-cognition and hippocampus. This result dovetails with previous research which has confirmed that hippocampal function (Ye et al., 2019; Zou & Kwok, 2022) and structure (Allen et al., 2017; Alkan et al., 2020) are highly correlated with meta-cognition. Further, given the essential role meta-cognition plays in other mentalising components according to interactive mentalising theory (Wu et al., 2020b), our finding supports the motivation to decipher the importance of the hippocampus during dynamic social interaction. Second, the variation of self-other mentalisation also showed greater representational similarity to individual variation in hippocampal MMS or rs-FC relative to amygdala MMS or rs-FC. There are two theoretical bases by which the hippocampus supports self-other mentalisation. Relational integration theory (O’Keefe & Nadel, 1978; Rubin et al., 2014) posits that the hippocampus buttresses various functions, such as memory, spatial reasoning and socialisation, through cognitive mapping, which denotes the capacity to organise and bind conceptual relationships (Tolman, 1948; Eichenbaum et al., 1999; Schiller et al., 2015). Constructive memory theory (Schacter, 2012) argues that how we retrieve and re-encode our memories can influence our sense of other individuals in the social milieu. Our finding is among the first to present additional evidence on the inter-individual level supporting the crucial role of the hippocampus in social cognition. Third, the variation of other-self mentalisation presented greater representational similarity to individual variation in amygdala MMS or rs-FC relative to hippocampal MMS or rs-FC. A plethora of literature has illustrated the importance of the amygdala in mentalising (Stone et al., 2003; Rice et al., 2014), especially in complex social interactions, such as trust or deception (Koscik & Tranel, 2011; Haas et al., 2015; Santos et al., 2016; Eskander et al., 2020). In concert with these studies, our result suggests a close link between other-self mentalisation, a higher-order mentalising component for the rich and complex social life, and the amygdala.

Finally, in favour of our **Hypothesis 3**, we found that high-rs-FC similarity boosted mentalising similarity in subject pairs with high-MMS similarity and lowered mentalising similarity for dyads with low-MMS similarity by leveraging the dyadic regression analysis (Fig. 7). As far as we know, no previous study has directly examined the interaction between brain morphometry and connectivity in predicting mentalising ability. Exploring such interaction is essential for elucidating the brain structure-function relationship, as recent studies demonstrate that functional connectivity links to the brain structure or structure connectome (Van Den Heuvel et al., 2009; Caparelli et al., 2017; Sorrentino et al., 2021; Levakov et al., 2021). Such a link, to some degree, has also been suggested to be abnormal in depression (Yun & Kim, 2021) and ASD (Hong et al., 2019). For instance, it might advance our understanding of ASD, as the complex nature of altered hippocampal structure-function interaction is evident in ASD (Banker et al., 2021). Our results thus provide additional evidence on this topic and offer further insights into social neuroscience that the brain structure differences may drive the levels of mentalising ability, which may be modulated by individual variations of rs-FC. Our study also makes two methodological contributions to the research community. First, it provides CPP-SD, a buttress against high-dimensional MMS data, thus paving the way for using MMS data in IS-RSA. The reason for the superiority of our proposed CPP-SD (i.e., computing patching and pooling operations-based surface distance) over other pipelines (Fig. 8) may be that patching and pooling operations over the surface could filter the noisy values, and computing surface distance based on these operations could offer a characterisation of surface-internal representations, which may further promote this effect. Second, our study provides a preliminary attempt (i.e., using grid search and subject-wise bootstrapping) to disentangle different parameter configurations in constructing IDMs. Although this approach may have a high computational load (depending on data type), it does not need to require specifying the specific parameter beforehand, e.g., the distance metric, which may represent some inappropriate assumptions about the structure of the brain-behaviour representational similarity (Finn et al., 2020) and thus lead researchers to underestimate this similarity. For example, different from the previous work (Chen et al., 2020), which suggests significant associations in IS-RSA only hold when using the multivariate representations, we found that the multivariate (item-wise dissimilarity) and univariate IMQ representations (the dissimilarity of composite score) are both essential to get the best performance (i.e., the highest correspondence between two IDMs) (Fig. 9). Our results indicate that both representations (constructed by the optimal parameter configuration obtained via grid search and bootstrapping) are theoretically appropriate, and each may capture different effects.

**Figure 9:**
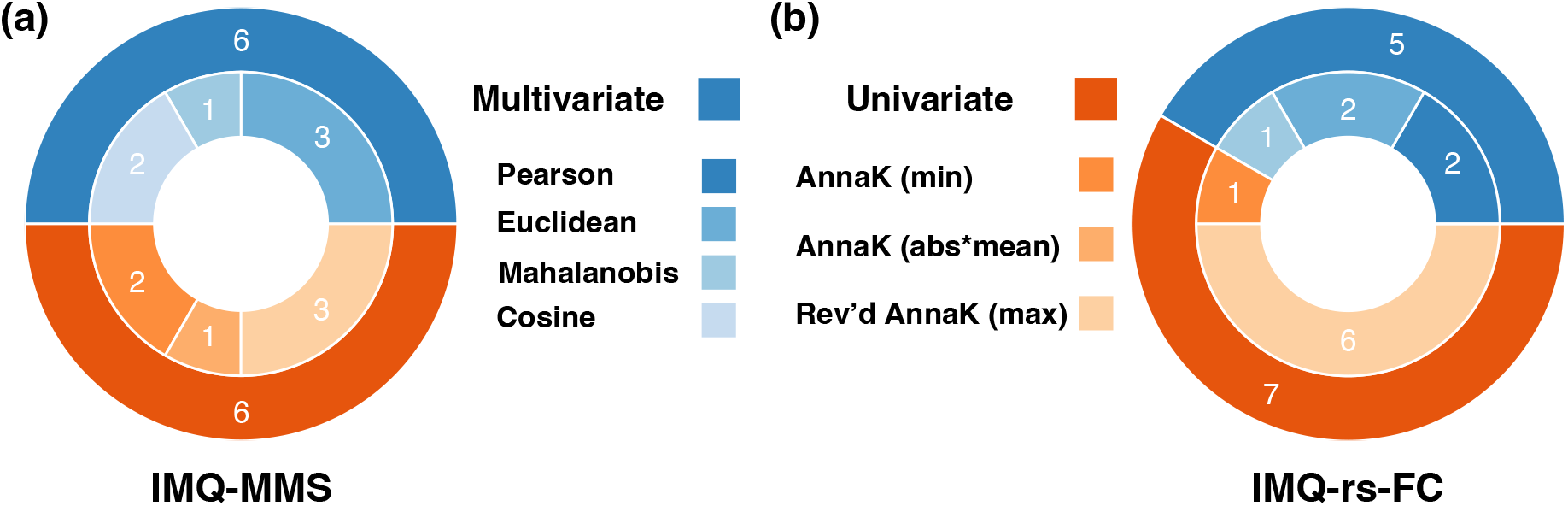
Comparison of winning IMQ representations. (a) In 12 cases for IMQ-MMS, the multivariate (i.e., item-wise dissimilarity) and univariate representations (i.e., the dissimilarity of composite score) are on par (6 vs 6) in determining the highest IDM correspondence; (b) Similarly, in 12 cases for IMQ-rs-FC, the multivariate and univariate representations have almost equal shares (5 vs 6) in the resulting highest IDM correspondence. The outer ring of the doughnut plot reflects representational indices (multivariate or univariate) of IMQ scores; the inner ring reflects the indices of the related distance metric (Pearson distance, Euclidean distance, Mahalanobis distance and cosine distance for construing the multivariate IMQ representations; Anna Karenina distances, including minimum distance and the product of the absolute and minimum distance, and reversed Anna Karenina distance, i.e., maximum distance, for construing the univariate IMQ representations). The figure was visualised using Matplotlib (Hunter, 2007).

The present study is only a first step toward capturing the neural underpinnings of interindividual variability in different aspects of social mentalising, and several limitations should be acknowledged. One potential limitation of our study, which probably has reduced the statistical power and increased the possibility of type II errors (Columb & Atkinson, 2016), is the relatively small number of participants (*N* = 24). Future work should try to replicate this study in larger samples and in a more nuanced way. Second, we did not unravel the potential lateralisation effects of the amygdala or hippocampus in our study. We caution against strong interpretations for lateralisation effects in our results because we had no prior specific hypotheses regarding lateralisation, and the lateralisation effects we observed are likely due to statistical thresholding effects. Third, we investigated how variation in rs-FC between the amygdala or hippocampus and other 115 brain regions mapped onto individual differences in the other two modalities. This means we could not reveal the contribution of rs-FC between the amygdala or hippocampus and specific brain regions or networks in the trinity. Future studies might uncover this by examining the rs-FC between the amygdala or hippocampus and some brain regions of interest, e.g., the temporoparietal junction (Biervoye et al., 2016; Bitsch et al., 2019), precuneus (Ye et al., 2019), cerebellum (Metoki et al., 2022) and angular gyrus (Zou & Kwok, 2022), or even the brain networks, e.g., the default mode network (Yeshurun et al., 2021).

## 5. Conclusion

The current work defines an integrative trinity framework that provides a testable basis for understanding individual differences in brain morphometry, connectivity and mentalising ability. Our study reveals the existence of a region-related specificity: the variation of SS and SO are more related to individual differences in hippocampal MMS and rs-FC, whereas the variation of OS shows a closer link with individual differences in amygdala MMS and rs-FC. Additionally, our data suggest that rs-FC gates the MMS predicted similarity in mentalising ability, revealing the intertwining role brain morphometry and connectivity play in social cognition. Broadly speaking, it is important to note that our mentalising ability to transition from an egocentric to an allocentric perspective, to appreciate, understand, predict and adapt to the intentions, thoughts and beliefs of others, makes us truly human. Ergo, to build a community with a shared future for mankind in today’s world, which is undergoing significant changes unseen in a century, understanding the neural underpinnings of individual differences in different mentalising components has never been more critical than today.

## Data and Code Availability Statement

The data and code used in this manuscript are available at https://github.com/andlab-um/trinity.

## Acknowledgement

This work is funded by the National Natural Science Foundation of China (U1736125), the Natural Science Foundation of Guangdong Province (2021A1515012509, 2019A1515111038), the Science and Technology Development Fund (FDCT) of Macau (0127/2020/A3), SRG of University of Macau (SRG2020-00027-ICI), Beijing Institute of Technology Research Fund Program for Young Scholars, the Fundamental Research Funds for the Central Universities (lzujbky-2021-kb26), the National Key Research and Development Program of China (2019YFA0706200), and the National Natural Science Foundation of China (61632014, 61627808). The authors would like to thank Mr Hao Yu, who provided general support in participant recruiting.

## Supplementary Material

**Table S1:**
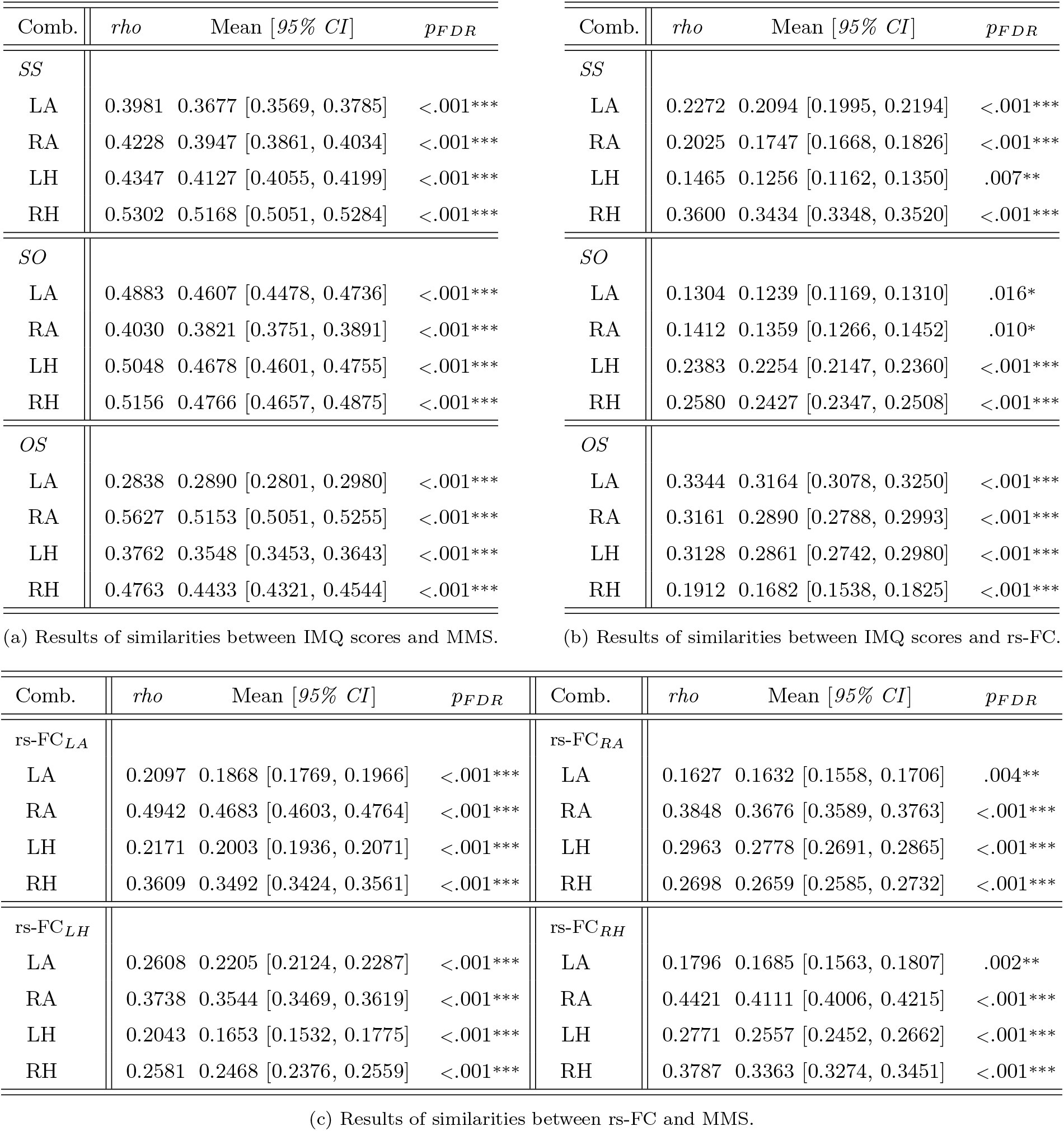
Detailed results of the trinity depicted in Fig. 6. ‘Comb.’ for combinations; ‘LA’ for the left amygdala; ‘RA’ for the right amygdala; ‘LH’ for the left hippocampus; ‘RH’ for the right hippocampus; ‘rho’ for Spearman’s rank correlation score between two IDMs. Mean values and 95% confidence intervals (numbers in brackets) were obtained by bootstrapping subjects. All the *p*-values were derived from the permutation test (10,000 iterations). We used FDR correction for multiple comparisons across all 40 statistical tests (12 for IMQ-MMS, 12 for IMQ-rs-FC and 16 for MMS-rs-FC). * *p* < .05, ** *p* < .01, *** *p* < .001.

## Footnotes

1 https://www.marxists.org/reference/archive/mao/selected-works/volume-1/mswv1_17.htm.

2 http://www.bic.mni.mcgill.ca/ServicesAtlases/HomePage

3 To better illustrate the possible effects, we entered similarities rather than dissimilarities between subject pairs (as we did in constructing IDMs) into the regression model. For each mentalising similarity measured by a different distance metric (we have ten different distance metrics for measuring mentalising similarity, as we mentioned before), we first chose related MMS and rs-FC similarity computed by the optimal parameter configuration (i.e., the configuration which offers the highest average Spearman correlation with the mentalising similarity from bootstrapping). Then we built ten regression models and chose the optimal model with the highest log-likelihood.

## References

Abraham, A., Pedregosa, F., Eickenberg, M., Gervais, P., Mueller, A., Kossaifi, J., Gramfort, A., Thirion, B., & Varoquaux, G. (2014). Machine learning for neuroimaging with scikitlearn. Frontiers in Neuroinformatics, 8, 1–10. doi:10.3389/fninf.2014.00014.

Adolphs, R. (2009). The social brain: Neural basis of social knowledge. Annual Review of Psychology, 60, 693–716. doi:10.1146/annurev.psych.60.110707.163514.

Alkan, E., Davies, G., Greenwood, K., & Evans, S. L. (2020). Brain Structural Correlates of Metacognition in First-Episode Psychosis. Schizophrenia Bulletin, 46, 552–561. doi:10.1093/schbul/sbz116.

Allen, J. G., & Fonagy, P. (2008). Handbook of Mentalization-Based Treatment. doi:10.1002/9780470712986.

Allen, M., Glen, J. C., Müllensiefen, D., Schwarzkopf, D. S., Fardo, F., Frank, D., Callaghan, M. F., & Rees, G. (2017). Metacognitive ability correlates with hippocampal and prefrontal microstructure. NeuroImage, 149, 415–423. doi:10.1016/j.neuroimage.2017.02.008.

Amodio, D. M., & Frith, C. D. (2006). Meeting of minds: The medial frontal cortex and social cognition. Nature Reviews Neuroscience, 7, 268–277. doi:10.1038/nrn1884.

Asen, E., & Fonagy, P. (2012). Mentalization-based Therapeutic Interventions for Families. Journal of Family Therapy, 34, 347–370. doi:10.1111/j.1467-6427.2011.00552.x.

Ashburner, J., & Friston, K. J. (2005). Unified segmentation. NeuroImage, 26, 839–851. doi:10.1016/j.neuroimage.2005.02.018.

Banker, S. M., Gu, X., Schiller, D., & Foss-Feig, J. H. (2021). Hippocampal contributions to social and cognitive deficits in autism spectrum disorder. Trends in Neurosciences, 44, 793–807. doi:10.1016/j.tins.2021.08.005.

Batista-García-Ramó, K., & Fernández-Verdecia, C. I. (2018). What we know about the brain structure-function relationship. Behavioral Sciences, 8. doi:10.3390/bs8040039.

Benjamini, Y., & Hochberg, Y. (1995). Controlling the false discovery rate: a practical and powerful approach to multiple testing. Journal of the Royal Statistical Society: Series B (Methodological), 57, 289–300. doi:10.1111/j.2517-6161.1995.tb02031.x.

Bickart, K. C., Dickerson, B. C., & Barrett, L. F. (2014). The amygdala as a hub in brain networks that support social life. Neuropsychologia, 63, 235–248. doi:10.1016/j.neuropsychologia.2014.08.013.

Biervoye, A., Dricot, L., Ivanoiu, A., & Samson, D. (2016). Impaired spontaneous belief inference following acquired damage to the left posterior temporoparietal junction. Social Cognitive and Affective Neuroscience, 11. doi:10.1093/scan/nsw076.

Bitsch, F., Berger, P., Nagels, A., Falkenberg, I., & Straube, B. (2019). Impaired right temporoparietal junction-hippocampus connectivity in schizophrenia and its relevance for generating representations of other minds. Schizophrenia Bulletin, 45, 934–945. doi:10.1093/schbul/sby132.

Bonner, M. F., & Epstein, R. A. (2018). Computational mechanisms underlying cortical responses to the affordance properties of visual scenes. PLoS Computational Biology, 14, 1–31. doi:10.1371/journal.pcbi.1006111.

Caparelli, E. C., Ross, T. J., Gu, H., Liang, X., Stein, E. A., & Yang, Y. (2017). Graph theory reveals amygdala modules consistent with its anatomical subdivisions. Scientific Reports, 7, 1–14. doi:10.1038/s41598-017-14613-4.

Chen, G., Shin, Y. W., Taylor, P. A., Glen, D. R., Reynolds, R. C., Israel, R. B., & Cox, R. W. (2016). Untangling the relatedness among correlations, part I: Nonparametric approaches to inter-subject correlation analysis at the group level. NeuroImage, 142, 248–259. doi:10.1016/j.neuroimage.2016.05.023.

Chen, G., Taylor, P. A., Shin, Y. W., Reynolds, R. C., & Cox, R. W. (2017). Untangling the relatedness among correlations, Part II: Inter-subject correlation group analysis through linear mixed-effects modeling. NeuroImage, 147, 825–840. doi:10.1016/j.neuroimage.2016.08.029.

Chen, P. H. A., Jolly, E., Cheong, J. H., & Chang, L. J. (2020). Intersubject representational similarity analysis reveals individual variations in affective experience when watching erotic movies. NeuroImage, 216, 116851. doi:10.1016/j.neuroimage.2020.116851.

Chua, E. F., Schacter, D. L., Rand-Giovannetti, E., & Sperling, R. A. (2006). Understanding metamemory: Neural correlates of the cognitive process and subjective level of confidence in recognition memory. NeuroImage, 29, 1150–1160. doi:10.1016/j.neuroimage.2005.09.058.

Collobert, R., Weston, J., Bottou, L., Karlen, M., Kavukcuoglu, K., & Kuksa, P. (2011). Natural language processing (almost) from scratch. Journal of Machine Learning Research, 12, 2493–2537.

Columb, M. O., & Atkinson, M. S. (2016). Statistical analysis: sample size and power estimations. BJA Education, 16, 159–161. doi:10.1093/bjaed/mkv034.

Davatzikos, C. (1996). Spatial normalization of 3D brain images using deformable models. Journal of computer assisted tomography, 20, 656–665. doi:10.1097/00004728-199607000-00031.

Dong, Q., Zhang, W., Stonnington, C. M., Wu, J., Gutman, B. A., Chen, K., Su, Y., Baxter, L. C., Thompson, P. M., Reiman, E. M., Caselli, R. J., & Wang, Y. (2020). Applying surface-based morphometry to study ventricular abnormalities of cognitively unimpaired subjects prior to clinically significant memory decline. NeuroImage: Clinical, 27. doi:10.1016/j.nicl.2020.102338.

Dong, Q., Zhang, W., Wu, J., Li, B., Schron, E. H., McMahon, T., Shi, J., Gutman, B. A., Chen, K., Baxter, L. C., Thompson, P. M., Reiman, E. M., Caselli, R. J., & Wang, Y. (2019). Applying surface-based hippocampal morphometry to study APOE-E4 allele dose effects in cognitively unimpaired subjects. NeuroImage: Clinical, 22. doi:10.1016/j.nicl.2019.101744.

Eichenbaum, H., & Cohen, N. J. (2014). Can We Reconcile the Declarative Memory and Spatial Navigation Views on Hippocampal Function? Neuron, 83, 764–770. doi:10.1016/j.neuron.2014.07.032.

Eichenbaum, H., Dudchenko, P., Wood, E., Shapiro, M., & Tanila, H. (1999). The hippocampus, memory, and place cells: Is it spatial memory or a memory space? Neuron, 23, 209–226. doi:10.1016/S0896-6273(00)80773-4.

Eickhoff, S. B., & Müller, V. I. (2015). Functional Connectivity. Brain Mapping: An Encyclopedic Reference, 2, 187–201. doi:10.1016/B978-0-12-397025-1.00212-8.

Eskander, E., Sanders, N., & Nam, C. S. (2020). Neural Correlates and Mechanisms of Trust. doi:10.1007/978-3-030-34784-0_22.

Feilong, M., Nastase, S. A., Guntupalli, J. S., & Haxby, J. V. (2018). Reliable individual differences in fine-grained cortical functional architecture. NeuroImage, 183, 375–386. doi:10.1016/j.neuroimage.2018.08.029.

Feurer, M., & Hutter, F. (2019). Hyperparameter Optimization. doi:10.1007/978-3-030-05318-5_1.

Finn, E. S., Glerean, E., Khojandi, A. Y., Nielson, D., Molfese, P. J., Handwerker, D. A., & Bandettini, P. A. (2020). Idiosynchrony: From shared responses to individual differences during naturalistic neuroimaging. NeuroImage, 215, 116828. doi:10.1016/j.neuroimage.2020.116828.

Finn, E. S., Shen, X., Scheinost, D., Rosenberg, M. D., Huang, J., Chun, M. M., Papademetris, X., & Constable, R. T. (2015). Functional connectome fingerprinting: Identifying individuals using patterns of brain connectivity. Nature Neuroscience, 18, 1664–1671. doi:10.1038/nn.4135.

Friston, K. J., Williams, S., Howard, R., Frackowiak, R. S. J., & Turner, R. (1996). Movement-Related effects in fMRI time-series. Magnetic Resonance in Medicine, 35, 346–355. doi:10.1002/mrm.1910350312.

Frith, C. D., & Frith, U. (2005). Theory of mind. Current biology, 15, 644–645. doi:10.1016/j.cub.2005.08.041.

Frith, C. D., & Frith, U. (2006). The Neural Basis of Mentalizing. Neuron, 50, 531–534. doi:10.1016/j.neuron.2006.05.001.

Frith, U., & Frith, C. (2001). The biological basis of social interaction. Current Directions in Psychological Science, 10, 151–155. doi:10.1111/1467-8721.00137.

Haas, B. W., Ishak, A., Anderson, I. W., & Filkowski, M. M. (2015). The tendency to trust is reflected in human brain structure. NeuroImage, 107, 175–181. doi:10.1016/j.neuroimage.2014.11.060.

Hong, S. J., de Wael, R. V., Bethlehem, R. A., Lariviere, S., Paquola, C., Valk, S. L., Milham, M. P., Di Martino, A., Margulies, D. S., Smallwood, J., & Bernhardt, B. C. (2019). Atypical functional connectome hierarchy in autism. Nature Communications, 10, 1–13. doi:10.1038/s41467-019-08944-1.

Hunter, J. D. (2007). Matplotlib: A 2D graphics environment. Computing in Science and Engineering, 9, 90–95. doi:10.1109/MCSE.2007.55.

Hyatt, C. J., Calhoun, V. D., Pittman, B., Corbera, S., Bell, M. D., Rabany, L., Pelphrey, K., Pearlson, G. D., & Assaf, M. (2020). Default mode network modulation by mentalizing in young adults with autism spectrum disorder or schizophrenia. NeuroImage: Clinical, 27, 102343. doi:10.1016/j.nicl.2020.102343.

Jolly, E. (2018). Pymer4: Connecting R and Python for Linear Mixed Modeling. Journal of Open Source Software, 3, 862. doi:10.21105/joss.00862.

Kaniuth, P., & Hebart, M. N. (2022). Feature-reweighted representational similarity analysis: A method for improving the fit between computational models, brains, and behavior. NeuroImage, 257, 119294. doi:10.1016/j.neuroimage.2022.119294.

Kerr, N., Dunbar, R. I., & Bentall, R. P. (2003). Theory of mind deficits in bipolar affective disorder. Journal of Affective Disorders, 73, 253–259. doi:10.1016/S0165-0327(02)00008-3.

Koscik, T. R., & Tranel, D. (2011). The human amygdala is necessary for developing and expressing normal interpersonal trust. Neuropsychologia, 49, 602–611. doi:10.1016/j.neuropsychologia.2010.09.023.

Koster-Hale, J., & Saxe, R. (2013). Functional neuroimaging of theory of mind. In Understanding Other Minds. doi:10.1093/acprof:oso/9780199692972.003.0009.

Kriegeskorte, N., & Kievit, R. A. (2013). Representational geometry: Integrating cognition, computation, and the brain. Trends in Cognitive Sciences, 17, 401–412. doi:10.1016/j.tics.2013.06.007.

Kriegeskorte, N., Mur, M., & Bandettini, P. (2008). Representational similarity analysis - connecting the branches of systems neuroscience. Frontiers in Systems Neuroscience, 2, 1–28. doi:10.3389/neuro.06.004.2008.

Kusner, M. J., Sun, Y., Kolkin, N. I., & Weinberger, K. Q. (2015). From Word Embeddings To Document Distances. In International conference on machine learning (pp. 957–966). volume 37.

Laurita, A. C., & Nathan Spreng, R. (2017). The hippocampus and social cognition. In The Hippocampus from Cells to Systems (pp. 537–558). doi:10.1007/978-3-319-50406-3_17.

Levakov, G., Faskowitz, J., Avidan, G., & Sporns, O. (2021). Mapping individual differences across brain network structure to function and behavior with connectome embedding. NeuroImage, 242, 118469. doi:10.1016/j.neuroimage.2021.118469.

Lu, Z., & Ku, Y. (2020). NeuroRA: A Python Toolbox of Representational Analysis From Multi-Modal Neural Data. Frontiers in Neuroinformatics, 14, 1–15. doi:10.3389/fninf.2020.563669.

MacLean, P. D. (1990). The triune brain in evolution: Role in paleocerebral functions.

Metoki, A., Wang, Y., & Olson, I. R. (2022). The Social Cerebellum: A Large-Scale Investigation of Functional and Structural Specificity and Connectivity. Cerebral Cortex, 32, 987–1003. doi:10.1093/cercor/bhab260.

Montagrin, A., Saiote, C., & Schiller, D. (2018). The social hippocampus. Hippocampus, 28, 672–679. doi:10.1002/hipo.22797.

Moritz, S., Gläscher, J., Sommer, T., Büchel, C., & Braus, D. F. (2006). Neural correlates of memory confidence. NeuroImage, 33, 1188–1193. doi:10.1016/j.neuroimage.2006.08.003.

Nguyen, M., Vanderwal, T., & Hasson, U. (2019). Shared understanding of narratives is correlated with shared neural responses. NeuroImage, 184, 161–170. doi:10.1016/j.neuroimage.2018.09.010.

Nummenmaa, L., Glerean, E., Viinikainen, M., Jääskeläinen, I. P., Hari, R., & Sams, M. (2012). Emotions promote social interaction by synchronizing brain activity across individuals. Proceedings of the National Academy of Sciences of the United States of America, 109, 9599–9604. doi:10.1073/pnas.1206095109.

O’Keefe, J., & Nadel, L. (1978). The hippocampus as a cognitive map.

Pang, L., Li, H., Liu, Q., Luo, Y.-J., Mobbs, D., & Wu, H. (2022). Resting-state functional connectivity of social brain regions predicts motivated dishonesty. NeuroImage, 256, 119253. doi:10.1016/j.neuroimage.2022.119253.

Parkinson, C., Kleinbaum, A. M., & Wheatley, T. (2018). Similar neural responses predict friendship. Nature Communications, 9. doi:10.1038/s41467-017-02722-7.

Pezzulo, G., Zorzi, M., & Corbetta, M. (2021). The secret life of predictive brains: what’s spontaneous activity for? Trends in Cognitive Sciences, 25, 730–743. doi:10.1016/j.tics.2021.05.007.

Pizer, S. M., Fritsch, D. S., Yushkevich, P. A., Johnson, V. E., & Chaney, E. L. (1999). Segmentation, registration, and measurement of shape variation via image object shape. IEEE Transactions on Medical Imaging, 18, 851–865. doi:10.1109/42.811263.

Popal, H., Wang, Y., & Olson, I. R. (2019). A Guide to Representational Similarity Analysis for Social Neuroscience. Social Cognitive and Affective Neuroscience, 14, 1243–1253. doi:10.1093/scan/nsz099.

Ren, Y., Nguyen, V. T., Sonkusare, S., Lv, J., Pang, T., Guo, L., Eickhoff, S. B., Breakspear, M., & Guo, C. C. (2018). Effective connectivity of the anterior hippocampus predicts recollection confidence during natural memory retrieval. Nature Communications, 9. doi:10.1038/s41467-018-07325-4.

Rice, K., Viscomi, B., Riggins, T., & Redcay, E. (2014). Amygdala volume linked to individual differences in mental state inference in early childhood and adulthood. Developmental Cognitive Neuroscience, 8, 153–163. doi:10.1016/j.dcn.2013.09.003.

Richell, R. A., Mitchell, D. G., Newman, C., Leonard, A., Baron-Cohen, S., & Blair, R. J. (2003). Theory of mind and psychopathy: Can psychopathic individuals read the ‘language of the eyes’? Neuropsychologia, 41, 523–526. doi:10.1016/S0028-3932(02)00175-6.

Rubin, R. D., Watson, P. D., Duff, M. C., & Cohen, N. J. (2014). The role of the hippocampus in flexible cognition and social behavior. Frontiers in Human Neuroscience, 8, 742. doi:10.3389/fnhum.2014.00742.

Santos, S., Almeida, I., Oliveiros, B., & Castelo-Branco, M. (2016). The role of the amygdala in facial trustworthiness processing: A systematic review and meta-analyses of fMRI studies. PLoS ONE, 11, 1–28. doi:10.1371/journal.pone.0167276.

Saxe, R., & Kanwisher, N. (2003). People thinking about thinking people: The role of the temporo-parietal junction in ”theory of mind”. NeuroImage, 19, 1835–1842. doi:10.1016/S1053-8119(03)00230-1.

Schaafsma, S. M., Pfaff, D. W., Spunt, R. P., & Adolphs, R. (2015). Deconstructing and reconstructing theory of mind. Trends in Cognitive Sciences, 19, 65–72. doi:10.1016/j.tics.2014.11.007.

Schacter, D. L. (2012). Adaptive constructive processes and the future of memory. American Psychologist, 67. doi:10.1037/a0029869.

Schafer, M., & Schiller, D. (2018). Navigating Social Space. Neuron, 100, 476–489. doi:10.1016/j.neuron.2018.10.006.

Schiller, D., Eichenbaum, H., Buffalo, E. A., Davachi, L., Foster, D. J., Leutgeb, S., & Ranganath, C. (2015). Memory and space: Towards an understanding of the cognitive map. Journal of Neuroscience, 35, 13904–13911. doi:10.1523/JNEUROSCI.2618-15.2015.

Schurz, M., Radua, J., Aichhorn, M., Richlan, F., & Perner, J. (2014). Fractionating theory of mind: A meta-analysis of functional brain imaging studies. Neuroscience and Biobehavioral Reviews, 42, 9–34. doi:10.1016/j.neubiorev.2014.01.009.

Schuwerk, T., Kaltefleiter, L. J., Au, J. Q., Hoesl, A., & Stachl, C. (2019). Enter the Wild: Autistic Traits and Their Relationship to Mentalizing and Social Interaction in Everyday Life. Journal of Autism and Developmental Disorders, 49, 4193–4208. doi:10.1007/s10803-019-04134-6.

Shepard, R. N., & Chipman, S. (1970). Second-order isomorphism of internal representations: Shapes of states. Cognitive Psychology, 1, 1–17. doi:10.1016/0010-0285(70)90002-2.

Shi, J., Thompson, P. M., Gutman, B., & Wang, Y. (2013). Surface fluid registration of conformal representation: Application to detect disease burden and genetic influence on hippocampus. NeuroImage, 78, 111–134. doi:10.1016/j.neuroimage.2013.04.018.

Siegal, M., & Varley, R. (2002). Neural systems involved in ‘theory of mind’. Nature Reviews Neuroscience, 3, 463–471. doi:10.1038/nrn844.

Snowden, J., Gibbons, Z., Blackshaw, A., Doubleday, E., Thompson, J., Craufurd, D., Foster, J., Happé, F., & Neary, D. (2003). Social cognition in frontotemporal dementia and Huntington’s disease. Neuropsychologia, 41, 688–701. doi:10.1016/S0028-3932(02)00221-X.

Sorrentino, P., Seguin, C., Rucco, R., Liparoti, M., Lopez, E. T., Bonavita, S., Quarantelli, M., Sorrentino, G., Jirsa, V., & Zalesky, A. (2021). The structural connectome constrains fast brain dynamics. eLife, 10, 1–11. doi:10.7554/eLife.67400.

Stone, V. E., Baron-Cohen, S., Calder, A., Keane, J., & Young, A. (2003). Acquired theory of mind impairments in individuals with bilateral amygdala lesions. Neuropsychologia, 41, 209–220. doi:10.1016/S0028-3932(02)00151-3.

Stuss, D. T., Gallup, G. G., & Alexander, M. P. (2001). The frontal lobes are necessary for ‘theory of mind’. Brain, 124, 279–286. doi:10.1093/brain/124.2.279.

Tavares, R. M., Mendelsohn, A., Grossman, Y., Williams, C. H., Shapiro, M., Trope, Y., & Schiller, D. (2015). A Map for Social Navigation in the Human Brain. Neuron, 87, 231–243. doi:10.1016/j.neuron.2015.06.011.

Thompson, P. M., Gledd, J. N., Woods, R. P., MacDonald, D., Evans, A. C., & Toga, A. W. (2000). Growth patterns in the developing brain detected by using continuum mechanical tensor maps. Nature, 404, 190–193. doi:10.1038/35004593.

Thompson, P. M., Hayashi, K. M., De Zubicaray, G. I., Janke, A. L., Rose, S. E., Semple, J., Hong, M. S., Herman, D. H., Gravano, D., Doddrell, D. M., & Toga, A. W. (2004). Mapping hippocampal and ventricular change in Alzheimer disease. NeuroImage, 22, 1754–1766. doi:10.1016/j.neuroimage.2004.03.040.

Tolman, E. C. (1948). Cognitive maps in rats and men. Psychological Review, 55, 189–208. doi:10.1037/h0061626.

Tzourio-Mazoyer, N., Landeau, B., Papathanassiou, D., Crivello, F., Etard, O., Delcroix, N., Mazoyer, B., & Joliot, M. (2002). Automated anatomical labeling of activations in SPM using a macroscopic anatomical parcellation of the MNI MRI single-subject brain. NeuroImage, 15, 273–289. doi:10.1006/nimg.2001.0978.

van Baar, J. M., Chang, L. J., & Sanfey, A. G. (2019). The computational and neural substrates of moral strategies in social decision-making. Nature Communications, 10. doi:10.1038/s41467-019-09161-6.

van Baar, J. M., Halpern, D. J., & FeldmanHall, O. (2021). Intolerance of uncertainty modulates brain-to-brain synchrony during politically polarized perception. Proceedings of the National Academy of Sciences of the United States of America, 118, 1–9. doi:10.1073/pnas.2022491118.

Van Den Heuvel, M. P., Mandl, R. C., Kahn, R. S., & Hulshoff Pol, H. E. (2009). Functionally linked resting-state networks reflect the underlying structural connectivity architecture of the human brain. Human Brain Mapping, 30, 3127–3141. doi:10.1002/hbm.20737.

Wang, J. X., Cohen, N. J., & Voss, J. L. (2015). Covert rapid action-memory simulation (CRAMS): A hypothesis of hippocampal-prefrontal interactions for adaptive behavior. Neurobiology of Learning and Memory, 117, 22–33. doi:10.1016/j.nlm.2014.04.003.

Wang, Y., Metoki, A., Xia, Y., Zang, Y., He, Y., & Olson, I. R. (2021). A large-scale structural and functional connectome of social mentalizing. NeuroImage, 236, 118115. doi:10.1016/j.neuroimage.2021.118115.

Wang, Y., Song, Y., Rajagopalan, P., An, T., Liu, K., Chou, Y. Y., Gutman, B., Toga, A. W., & Thompson, P. M. (2011). Surface-based TBM boosts power to detect disease effects on the brain: An N=804 ADNI study. NeuroImage, 56, 1993–2010. doi:10.1016/j.neuroimage.2011.03.040.

Wang, Y., Zhang, J., Gutman, B., Chan, T. F., Becker, J. T., Aizenstein, H. J., Lopez, O. L., Tamburo, R. J., Toga, A. W., & Thompson, P. M. (2010). Multivariate tensor-based morphometry on surfaces: Application to mapping ventricular abnormalities in HIV/AIDS. NeuroImage, 49, 2141–2157. doi:10.1016/j.neuroimage.2009.10.086.

Washburn, D., Wilson, G., Roes, M., Rnic, K., & Harkness, K. L. (2016). Theory of mind in social anxiety disorder, depression, and comorbid conditions. Journal of Anxiety Disorders, 37, 71–77. doi:10.1016/j.janxdis.2015.11.004.

Waskom, M. (2021). Seaborn: Statistical Data Visualization. Journal of Open Source Software, 6, 3021. doi:10.21105/joss.03021.

Worker, A., Dima, D., Combes, A., Crum, W. R., Streffer, J., Einstein, S., Mehta, M. A., Barker, G. J., Williams, S. C. R., & O’daly, O. (2018). Test–retest reliability and longitudinal analysis of automated hippocampal subregion volumes in healthy ageing and alzheimer’s disease populations. Human Brain Mapping, 39, 1743–1754. doi:10.1002/hbm.23948.

Wu, H., Feng, C., Lu, X., Liu, X., & Liu, Q. (2020a). Oxytocin effects on the resting-state mentalizing brain network. Brain Imaging and Behavior, 14. doi:10.1007/s11682-019-00205-5.

Wu, H., Fung, B. J., & Mobbs, D. (2022). Mentalizing During Social Interaction: The Development and Validation of the Interactive Mentalizing Questionnaire. Frontiers in Psychology, 12. doi:10.3389/fpsyg.2021.791835.

Wu, H., Liu, X., Hagan, C. C., & Mobbs, D. (2020b). Mentalizing during social InterAction: A four component model. Cortex, 126, 242–252. doi:10.1016/j.cortex.2019.12.031.

Wu, J., Dong, Q., Gui, J., Zhang, J., Su, Y., Chen, K., Thompson, P. M., Caselli, R. J., Reiman, E. M., Ye, J., & Wang, Y. (2021). Predicting Brain Amyloid Using Multivariate Morphometry Statistics, Sparse Coding, and Correntropy: Validation in 1,101 Individuals From the ADNI and OASIS Databases. Frontiers in Neuroscience, 15, 1–17. doi:10.3389/fnins.2021.669595.

Xia, M., Wang, J., & He, Y. (2013). BrainNet Viewer: A Network Visualization Tool for Human Brain Connectomics. PLoS ONE, 8, e68910. doi:10.1371/journal.pone.0068910.

Yan, C.-G., Wang, X.-D., Zuo, X.-N., & Zang, Y.-F. (2016). DPABI: Data Processing & Analysis for (Resting-State) Brain Imaging. Neuroinformatics, 14, 339–351. doi:10.1007/s12021-016-9299-4.

Yao, Z., Fu, Y., Wu, J., Zhang, W., Yu, Y., Zhang, Z., Wu, X., Wang, Y., & Hu, B. (2020). Morphological changes in subregions of hippocampus and amygdala in major depressive disorder patients. Brain Imaging and Behavior, 14, 653–667. doi:10.1007/s11682-018-0003-1.

Ye, Q., Zou, F., Dayan, M., Lau, H., Hu, Y., & Kwok, S. C. (2019). Individual susceptibility to TMS affirms the precuneal role in meta-memory upon recollection. Brain Structure and Function, 224, 2407–2419. doi:10.1007/s00429-019-01909-6.

Yeshurun, Y., Nguyen, M., & Hasson, U. (2021). The default mode network: where the idiosyncratic self meets the shared social world. Nature Reviews Neuroscience, 22, 181–192. doi:10.1038/s41583-020-00420-w.

Yokoi, S., Takahashi, R., Akama, R., Suzuki, J., & Inui, K. (2020). Word rotator’s distance. In Proceedings of the 2020 Conference on Empirical Methods in Natural Language Processing (pp. 2944–2960). doi:10.18653/v1/2020.emnlp-main.236.

Yun, J. Y., & Kim, Y. K. (2021). Graph theory approach for the structural-functional brain connectome of depression. Progress in Neuro-Psychopharmacology and Biological Psychiatry, 111, 110401. doi:10.1016/j.pnpbp.2021.110401.

Zou, F., & Kwok, S. C. (2022). Distinct generation of subjective vividness and confidence during naturalistic memory retrieval in angular gyrus. Journal of Cognitive Neuroscience, (pp. 1–13). doi:10.1162/jocn_a_01838.

